# Molecular Regulation and Physiological Role of GOLPH3-mediated Golgi retention

**DOI:** 10.1101/2025.06.26.661665

**Authors:** Anastasia Theodoropoulou, Anita Nasrallah, Luciano A. Abriata, Laurence Abrami, Francesco Talotta, Maria J. Marcaida, Muhammad U Anwar, Ondrej Kováč, Sergey Y. Vakhrushev, Alejandro Calleja, Sylvia Ho, Antonino Asaro, Francisco S Mesquita, Leila Alieh, Charlotte Gehin, Lucie Bracq, Irmak Kaysudu, Nika Gorsek, Sarah Vacle, Arthur Samurkas, Alessio Prunotto, Miroslav Machala, Olaia Naveiras, Katrine T Schjoldager, F. Gisou van der Goot, Matteo Dal Peraro, Giovanni D’Angelo

**Affiliations:** Laboratory for Biomolecular Modeling, Institute of Bioengineering, School of Life Sciences, École Polytechnique Fédérale de Lausanne (EPFL), Lausanne, Switzerland; Laboratory of Lipid Cell Biology, Institute of Bioengineering and Global Health Institute, School of Life Sciences, École Polytechnique Fédérale de Lausanne (EPFL), Lausanne, Switzerland; Global Health Institute, School of Life Sciences, EPFL, Lausanne, Switzerland; Institute of Genetics and Biophysics, National Research Council, Naples, Italy; Veterinary Research Institute, Department of Pharmacology and Toxicology, 62100 Brno, Czech Republic; Copenhagen Center for Glycomics, Departments of Cellular and Molecular Medicine, University of Copenhagen Copenhagen, Denmark; Laboratory of Regenerative Hematopoiesis, Department of Biomedical Sciences, Faculty of Biology and Medicine, University of Lausanne, Lausanne, Switzerland; Hematology Service, Departments of Oncology and Laboratory Medicine, Lausanne University Hospital (CHUV), Lausanne, Switzerland

## Abstract

The Golgi complex serves as the central hub of the biosynthetic pathway, where anterograde and retrograde trafficking converge. How cargo and Golgi-resident proteins traverse this organelle has long been debated. Recent studies have identified a molecular machinery that sorts resident proteins into retrograde-directed COPI vesicles during cisternal maturation. Golgi phosphoprotein 3 (GOLPH3) is a key component of this system; however, its physiological relevance and regulatory mechanisms remain poorly defined. Here, we show that GOLPH3 depletion in mice disrupts both protein and lipid glycosylation, causes partially penetrant embryonic lethality, and severely impairs growth and bone mineralization. At the molecular level, we find that GOLPH3 is regulated by functionally antagonistic S-acylation events that control the topology of its membrane association. To mediate retrograde trafficking of Golgi-resident glycosyltransferases, GOLPH3 must bind their cytosolic tails. This occurs via a negatively charged surface region, which is correctly oriented only in one of the S-acylated GOLPH3 conformations. Together, these findings reveal a lipid-mediated regulatory mechanism for intra-Golgi trafficking and establish the critical role of GOLPH3 in vertebrate development.

## Introduction

Proteins and lipids synthesized in the endoplasmic reticulum (ER) are transported to the Golgi complex, where resident enzymes modify them prior to their delivery to final cellular destinations (*1*). The most prominent of these modifications is glycosylation, catalyzed by approximately 200 glycosyltransferases that collectively define the mammalian glycome (*2*). Unlike protein or nucleic acid synthesis, glycan assembly proceeds without relying on a nucleic acid template (*3*). Nevertheless, it is both highly reproducible and dynamically responsive. The mechanisms that enable this balance of fidelity and flexibility remain only partially understood.

The Golgi apparatus is thought to form from ER-derived carriers that fuse at its *cis* face to generate new *cisternae*. These *cisternae* then mature into medial and *trans* compartments through the sequential remodeling of their glycosyltransferase content, while cargo remains enclosed within the lumen or membrane (*4*). Glycosyltransferases are thus continuously recycled within the maturing Golgi stack to support the ordered assembly of glycans on proteins and lipids (*5*).

Glycosyltransferase recycling depends on retrograde retrieval via Coat Protein complex 1 (COPI)-coated vesicles, which return enzymes to earlier Golgi compartments. Sorting into these vesicles requires either direct interaction with COPI subunits (*6*) or indirect recruitment through adaptor proteins that link enzymes to the COPI machinery (*7*, *8*). Although the molecular mechanisms of this sorting process are well characterized, its physiological regulation and broader impact remain poorly defined. In particular, it is unclear how this trafficking system adapts to modulate the glycome in response to cellular cues.

To address this gap, we investigated Golgi phosphoprotein 3 (GOLPH3), the first identified COPI adaptor (*9–11*). GOLPH3 is a peripheral membrane protein that localizes to the *trans*-Golgi by binding phosphatidylinositol-4-phosphate (PtdIns(4)*P*) (*12*, *13*). Adjacent to its PtdIns(4)*P*-binding site is a hydrophobic β-hairpin that inserts into the membrane and promotes curvature (*12*, *14*, *15*). Once membrane-bound, GOLPH3 interacts with COPI coatomer via an N-terminal basic motif (*9*), and with a subset of glycosyltransferases, facilitating their retrograde transport and protecting them from lysosomal degradation (*10*, *11*). GOLPH3 preferentially recognizes a basic residue–enriched motif in the cytosolic tails of glycosyltransferases (*9–11*), which are typically type II transmembrane proteins with luminal catalytic domains (*16*).

GOLPH3 clients span multiple glycosylation pathways, including glycosphingolipid biosynthesis (*17*), N-linked and mucin-type O-linked glycosylation, proteoglycan synthesis, O-mannosylation, tyrosine sulfation, and nucleotide hydrolysis (*11*). GOLPH3 also maintains lysosomal function by stabilizing LYSET, a key component of the mannose-6-phosphate tagging system (*18*). Accordingly, perturbing GOLPH3 expression alters both protein and lipid glycosylation in cultured cells (*11*, *19*).

Among its clients, GOLPH3 critically regulates lactosylceramide synthase (LCS, also known as B4GALT5), a *trans*-Golgi enzyme involved in glycosphingolipid biosynthesis (*10*). GOLPH3 anchors LCS at the Golgi, shielding it from proteolytic degradation (*17*). LCS receives its substrate, glucosylceramide, from the lipid transfer protein FAPP2, another PtdIns(4)*P* effector, and catalyzes the production of complex glycosphingolipids (*20*, *21*). By stabilizing LCS, GOLPH3 plays a key role in lipid remodeling at the Golgi. GOLPH3 also regulates mucin-type O-glycosylation by retaining GALNT family enzymes, which initiate glycan attachment to serine and threonine residues (*22*). Loss of GOLPH3 leads to degradation or mislocalization of these enzymes, impairing glycoprotein processing, disrupting glycan biosynthesis, and compromising extracellular matrix organization (*11*).

In this study, we examined the physiological consequences of GOLPH3 dysfunction in a mammalian model and uncovered an essential role in development. We next investigated how cells regulate GOLPH3 membrane association and client interactions. Our results reveal a novel regulatory mechanism involving antagonistic S-acylation events that establish distinct membrane topologies, either enabling or inhibiting client engagement. These findings demonstrate that GOLPH3-mediated retention of glycosyltransferases can be dynamically modulated by a previously unrecognized mechanism of intra-Golgi trafficking control. This work raises new questions about how acute and post-transcriptional regulatory pathways converge to reshape the glycome, and guide organismal development.

## Results

### GOLPH3 loss of function induces glycosylation defects in vivo

While GOLPH3 overexpression has been linked to solid tumor progression (*23*), the physiological impact of GOLPH3 loss of function *in vivo* remains largely unknown. To address this, we investigated the systemic effects of GOLPH3 deficiency in mice (**Figure 1A**). We first evaluated GOLPH3 expression at the mRNA and protein levels across multiple tissues in male and female mice at 15 and 41 weeks of age. GOLPH3 was ubiquitously expressed with minimal variation across age or sex (**Figure S1A, B**), consistent with single-cell RNA-seq data from the Tabula Muris atlas (*24*), which reports broad GOLPH3 expression in 20 mouse tissues (**Figure S1C**). Notably, GOLPH3 has a paralog, GOLPH3L, with overlapping function but more restricted expression, primarily in highly glycosylating cells such as goblet cells in the large intestine (**Figure S1C**).

**Figure 1.**
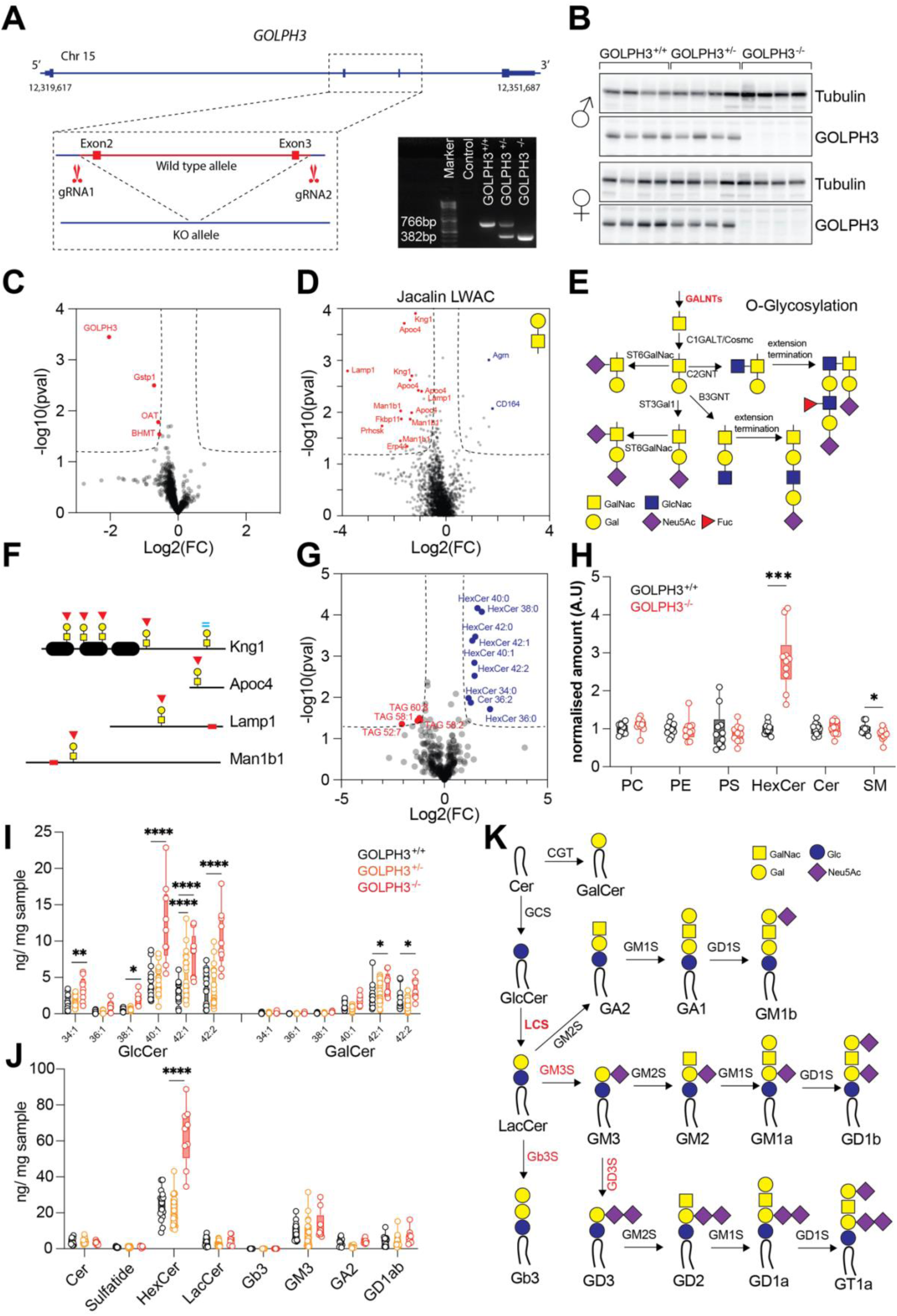
GOLPH3 loss disrupts glycosylation and lipid metabolism in mouse liver. A. Schematic of the CRISPR/Cas9-mediated strategy used to generate GOLPH3 knockout (GOLPH3^-/-^) mice by deleting exons 2 and 3. The diagnostic PCR (right) confirms wild-type (766 bp) and KO (382 bp) alleles in genomic DNA from mice of the indicated genotypes. **B.** Western blot analysis of liver lysates from male (top) and female (bottom) mice with the indicated genotypes. GOLPH3 and Tubulin (loading control) were detected with specific antibodies, confirming complete loss of GOLPH3 protein in knockout livers. **C.** Volcano plot showing differential protein abundance in livers from GOLPH3^+/+^ vs GOLPH3^-/-^ mice (n = 3) by TMT-based quantitative proteomics. GOLPH3 is the most significantly downregulated protein. **D.** Volcano plot showing changes in O-glycopeptide abundance from the same samples as in (C), enriched using jacalin-based lectin weak affinity chromatography (LWAC) and analyzed by TMT-MS. Several known GALNT2 substrates (e.g., Kng1, Apoc4, Lamp1) show reduced glycosylation in GOLPH3^-/-^ livers. **E.** Diagram of the mucin-type O-glycosylation pathway, illustrating key enzymes and intermediates. GALNTs initiate glycosylation by transferring GalNAc to Ser/Thr residues. **F.** Schematic showing O-glycosylation sites in glycoproteins (e.g., Kng1, Apoc4, Lamp1) with decreased glycopeptide abundance in GOLPH3^-/-^ livers. Red arrowheads indicate a decrease; cyan equality symbols indicate no change. **G.** Volcano plot depicting differential lipid species abundance between GOLPH3^+/+^ and GOLPH3^-/-^ livers (n = 10). Hexosylceramides (HexCer, blue) are elevated, while triglycerides (TAGs, red) are reduced in GOLPH3^-/-^ males. **H.** Box-and-whisker plots showing normalized abundance of major lipid classes. HexCer levels are significantly increased in the experimental group compared to controls (unpaired t-test, ***p < 0.001), while SM shows minor differences (n = 10 per group) **I.** Quantification of individual GlcCer and GalCer species via targeted LC-MS/MS across genotypes. GlcCer species are significantly elevated in GOLPH3^-/-^ livers; GalCer levels show minor increases (two-way ANOVA, *p < 0.05; **p < 0.001; ****p < 0.0001). **J.** Quantification of glycosphingolipid intermediates and products via LC-MS/MS. GlcCer accumulates in GOLPH3^-/-^ livers, with no significant change in downstream LacCer and gangliosides, consistent with impaired LCS activity (two-way ANOVA, * ****p < 0.0001). **K.** Schematic of the glycosphingolipid biosynthesis pathway. GOLPH3-dependent LCS converts GlcCer to LacCer, which serves as a precursor for downstream gangliosides and globosides. Accumulation of GlcCer in GOLPH3^-/-^ mice indicates reduced LCS function.

We generated GOLPH3 loss of function (GOLPH3^-/-^) mice via CRISPR/Cas-mediated deletion of exons 2 and 3 (**Figure 1A**, see methods). Western blotting on knockout livers confirmed loss of GOLPH3 protein (**Figure 1B**).

To examine the consequences for glycosylation, we focused on the liver, a key site of glycoprotein production. Tandem mass tag (TMT)-based quantitative proteomics identified few significant changes among >2,500 peptides in GOLPH3^-/-^ livers, with GOLPH3 itself being the most significantly downregulated protein (**Figure 1C**). This residual GOLPH3 detection in GOLPH3^-/-^ livers is likely linked to our gene targeting strategy (see methods). However, when the same samples were analyzed using lectin weak affinity chromatography (LWAC) followed by TMT-MS to enrich for O-glycopeptides, a distinct subset was markedly reduced. These included known substrates of the GOLPH3 client GALNT2, such as APOC4 and LAMP1 (*25*), suggesting impaired O-glycosylation due to client misregulation (**Figure 1D–F**).

We next performed untargeted LC-MS lipidomics on GOLPH3^-/-^ livers. Most lipid classes were unchanged, but hexosylceramide (HexCer) levels were consistently elevated in both male and female knockouts (**Figure 1G–H**). In contrast, triglyceride levels were selectively reduced in male GOLPH3^-/-^ livers (**Figure S1D**).

Because standard LC-MS cannot differentiate isobaric species like glucosylceramide (GlcCer) and galactosylceramide (GalCer), we employed targeted LC-MS/MS. This analysis revealed a significant increase in GlcCer, but not GalCer, in GOLPH3^-/-^ livers compared to controls (**Figure 1I**). Further sphingolipidomic profiling confirmed elevated GlcCer levels, while downstream metabolites of LCS activity remained unchanged (**Figure 1J**).

These findings indicate reduced LCS activity in the absence of GOLPH3, resulting in a metabolic bottleneck and GlcCer accumulation (**Figure 1K**). This phenotype parallels our earlier findings in HeLa cells, where GOLPH3 knockdown impaired LCS localization and stability (*17*). Together, our results demonstrate that GOLPH3 orchestrates glycosylation and lipid metabolism *in vivo*, likely by maintaining the proper localization and function of its enzyme clients.

### GOLPH3 loss of function results in partially penetrant embryonic lethality and bone defects

Strikingly, GOLPH3^+/-^ intercrosses yielded a significant deviation from Mendelian expectations. Among 552 offspring, only 77 (14%) were GOLPH3^-/-^, compared to 154 (28%) GOLPH3^+/+^ and 321 (58%) GOLPH3^+/-^ (χ², *p* < 0.0001), indicating substantial embryonic lethality in GOLPH3-null mice (**Figure 2A**).

**Figure 2:**
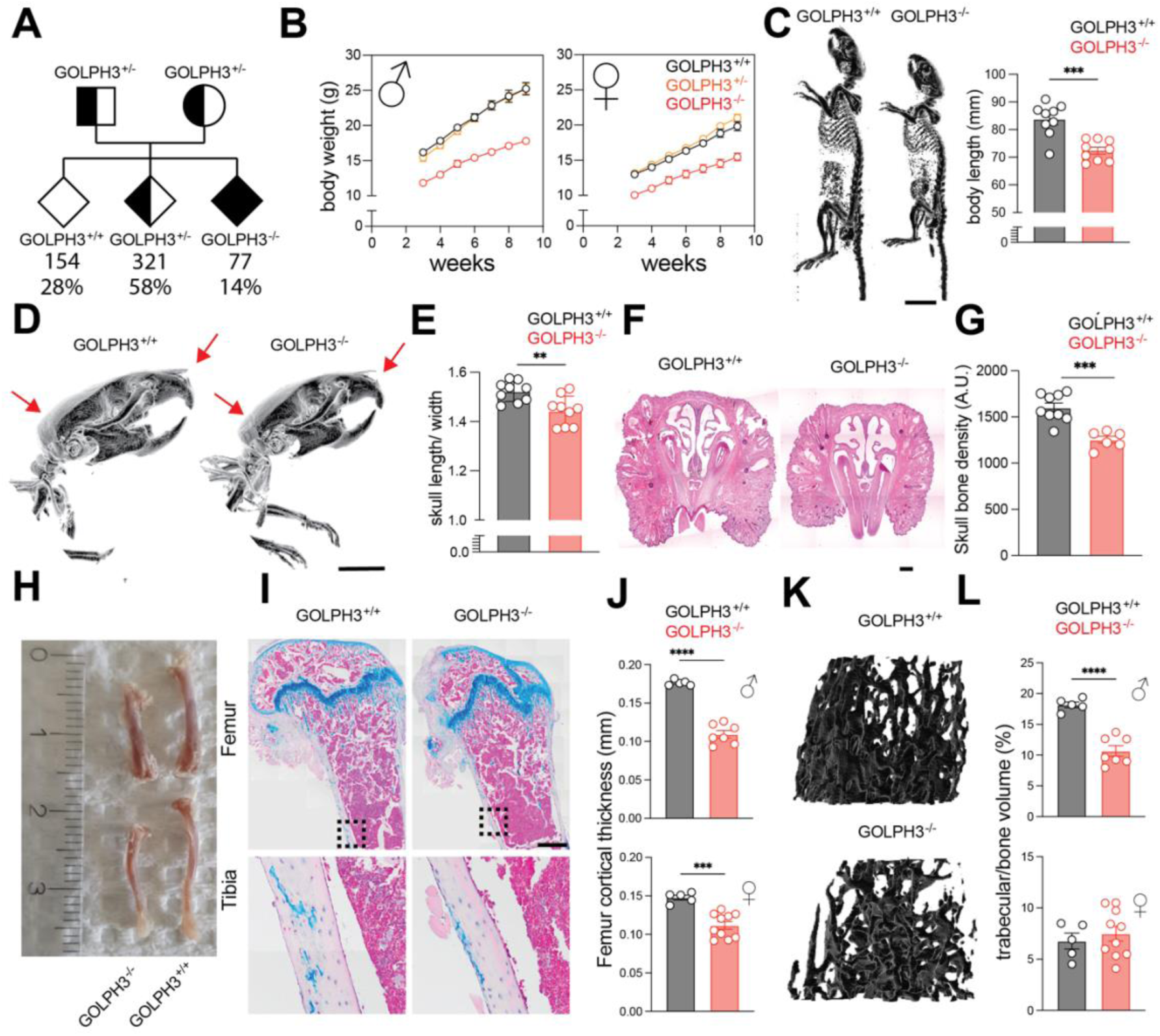
GOLPH3 loss of function leads to partially penetrant embryonic lethality and bone defects. A. Pedigree chart showing genotype frequencies among offspring from GOLPH3^+/-^ × GOLPH3^+/-^ crosses. **B.** Growth curves displaying body weight trajectories for male (left) and female (right) GOLPH3^+/+^, GOLPH3^+/-^, and GOLPH3^-/-^ mice between 3 and 9 weeks of age. Data are presented as mean ± SEM. **C.** Representative whole-body micro-computed tomography (µCT) scans of 9-week-old GOLPH3^+/+^ and GOLPH3^-/-^ mice (left). Quantification of body length is shown on the right. Data are mean ± SEM (unpaired t-test, ***p < 0.001). Bar = 10mm. **D.** Representative µCT scans of the skull from 9-week-old male GOLPH3^+/+^ and GOLPH3^-/-^ mice. Red arrows highlight craniofacial anomalies, including reduced snout length and diminished cranial bulging in GOLPH3^-/-^ mice. Bar = 5mm. **E.** Quantification of skull length-to-width ratio in 9-week-old GOLPH3^+/+^ and GOLPH3^-/-^ mice. Data are mean ± SEM (unpaired t-test, **p < 0.01). **F.** Representative hematoxylin and eosin (H&E)-stained coronal sections of the nasal cavity in 9-week-old GOLPH3^+/+^ and GOLPH3^-/-^ mice. Bar = 1mm. **G.** Quantification of skull bone density in GOLPH3^+/+^ and GOLPH3^-/-^ mice assessed by µCT. Data are mean ± SEM (unpaired t-test, ***p < 0.001). **H.** Representative photograph of femurs and tibias from 9-week-old GOLPH3^+/+^ and GOLPH3^-/-^ mice. **I.** Representative Alcian Blue-stained decalcified femur sections from 9-week-old GOLPH3^+/+^ and GOLPH3^-/-^ mice. Insets show magnified views of cortical bone regions. Bar = 500µm. **J.** µCT-based quantification of femoral cortical thickness in male (top) and female (bottom) GOLPH3^+/+^ and GOLPH3^-/-^ mice. Data are mean ± SEM (unpaired t-test, ***p < 0.001; ****p<0.0001). **K.** Representative 3D reconstructions of femoral trabecular bone from 9-week-old male GOLPH3^+/+^ and GOLPH3^-/-^ mice. Bar = 200µm **L.** Quantification of femoral trabecular bone volume fraction (%) in male (top) and female (bottom) mice. Data are mean ± SEM (unpaired t-test, ****p < 0.0001).

Surviving GOLPH3^-/-^ mice showed marked growth defects. Both males and females exhibited a 20–30% reduction in body weight between weeks 3 and 9 (**Figure 2B**), driven by losses in both fat and lean mass as shown by EchoMRI (**Figure S2A**). Body length was also significantly reduced (**Figure 2C**). While fasting glucose levels were lower in GOLPH3^-/-^ mice, insulin and glucose tolerance remained unchanged across genotypes (**Figure S2B, C**). Organ weights were generally reduced, but this effect was normalized when adjusted for total body weight, except in the brain, which showed a relatively increased mass in GOLPH3^-/-^ mice (**Figure S2D**).

Craniofacial and skeletal anomalies further revealed a role for GOLPH3 in bone development. GOLPH3^-/-^ mice had shorter snouts, reduced skull doming (**Figure 2D, E**), and diminished skull bone density on µCT, despite preserved nasal cavity structure (**Figure 2F, G**). Femoral and tibial lengths were ∼66% of controls in both sexes (**Figure 2H**, and **S2E**), indicating global skeletal stunting. Cortical bone thickness was also reduced, particularly in males, by 30% in the femur and 15% in the tibia, whereas females showed milder effects (**Figure 2I, J; Figure S2F**).

Trabecular bone defects exhibited sex-specific differences: male GOLPH3^-/-^ mice had significantly less dense trabeculae compared to controls, a phenotype not observed in females (**Figure 2K, L; Figure S2G**). Together, these findings demonstrate that GOLPH3 loss impairs growth and bone architecture, with more severe consequences in males. Notably, some skeletal features resemble those of GALNT2 knockout mice (*26*), linking GOLPH3-dependent Golgi trafficking to bone development via its glycosyltransferase clients.

### Regulated topology of GOLPH3 membrane association

The evidence above underscores the physiological importance of GOLPH3. This primed us to investigate its role as a Golgi adapter at the molecular level. Because client interaction depends on GOLPH3’s membrane association, we first examined its binding to the Golgi bilayer using coarse-grained (CG) molecular dynamics (MD) simulations.

We initialized GOLPH3 (PDB: 3KN1) at ∼60 Å from a membrane modeled with a lipid composition based on lipidomics of Golgi isolates, including 5% molar PtdIns(4)*P* (*27*) (see Methods) (**Figure 3A**). Simulations revealed rapid membrane association, with GOLPH3 binding after 0.63 ± 0.26 µs on average (**Figure S3A**).

**Figure 3.**
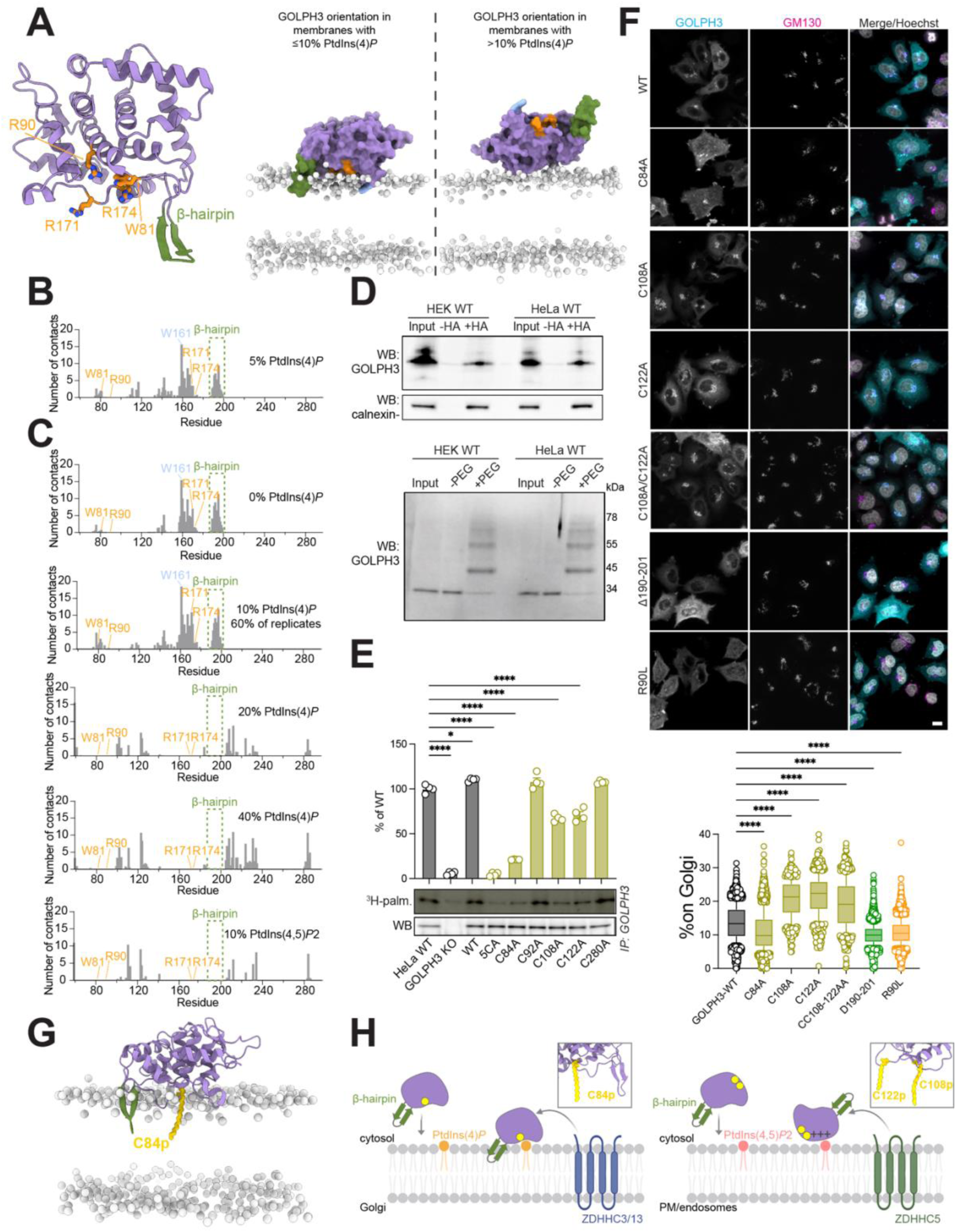
GOLPH3 membrane orientation. A. Structure of GOLPH3 (left) and snapshots from coarse-grained MD simulations with membranes containing 5% (middle) or 10% (right) PtdIns(4)P. At 5%, GOLPH3 (purple surface) interacts via its β-hairpin (green), PtdIns(4)*P*-binding residues (W81, R90, R171, R174; orange), and Trp161 (blue). At 10% and higher, binding is mediated by a positively charged surface. Lipid head groups are shown as grey spheres; other lipids and water are omitted for clarity. **B.** Average number of contacts between GOLPH3 residues and membranes with 5% PtdIns(4)P across MD trajectories. **C.** MD-based contact analysis of GOLPH3 with membranes containing 0%, 10% (detected in 60% of replicates), 20%, 40% PtdIns(4)*P*, or 10% PtdIns(4,5)*P*₂. **D.** S-acylation of endogenous GOLPH3 in HEK and HeLa cells. Palmitoylated proteins were detected by hydroxylamine treatment (+HA), followed by SDS-PAGE and anti-GOLPH3 immunoblotting. Calnexin was used as a loading control (Top). Stoichiometry of GOLPH3 palmitoylation determined via PEG-5 labeling and anti-GOLPH3 western blotting (Bottom). **E.** Palmitoylation analysis of GOLPH3 cysteine mutants. GOLPH3 KO HeLa cells were transfected with WT or mutant constructs, labeled with ³H-palmitic acid, and analyzed by autoradiography following immunoprecipitation. Quantification (mean ± SEM, n = 4) is relative to endogenous WT GOLPH3. **F.** Immunofluorescence of HeLa cells expressing WT or mutant GOLPH3, labeled with antibodies against GOLPH3, GM130 (Golgi), and Hoechst (nuclei). Scale bars = 10 μm (Top). Quantification of Golgi-to-cytosol GOLPH3 intensity ratio (Bottom). Whiskers show 2.5th–97.5th percentile; outliers as dots. One-way ANOVA vs. WT: ****p < 0.0001. Only transfected cells were analyzed (defined by GOLPH3 intensity > mean + 10 SD of non-transfected cells). **G.** MD model of S-acylated GOLPH3 (C84) interacting with membranes. **H.** Proposed model of GOLPH3 membrane recruitment, integrating electrostatic interactions (via PtdIns(4)*P*-binding residues or a positively charged surface), β-hairpin insertion, and S-acylation at C84, C108, and C122.

GOLPH3 consistently approached the membrane via a specific surface. In this orientation, the hydrophobic β-hairpin (residues 190–201) and W161 inserted into the bilayer, while key PtdIns(4)*P*-binding residues (W81, R90, R171, R174) (*12*, *13*) were positioned near the membrane (**Figure 3A, B**). This orientation aligns with models from OPM and AlphaFold3 (*28*, *29*) (**Figure S3B, C**), and with prior biochemical studies using purified GOLPH3 (*12*).

To assess how PtdIns(4)*P* influences membrane binding, we ran CG MD simulations with increasing concentrations of PtdIns(4)*P* (0–40% molar). GOLPH3 associated with membranes even in the absence of PtdIns(4)*P* (binding in 5 of 6 MD replicas), maintaining the same binding surface as at physiological PtdIns(4)*P* levels (5%) (**Figure 3C**).

Remarkably, at higher PtdIns(4)*P* levels (10%), GOLPH3 adopted an alternative orientation in ∼40% of cases, engaging the membrane via a completely distinct, positively charged surface located on the opposing side of the protein (**Figure 3A, C**). This flipped conformation became exclusive at 20% and 40% PtdIns(4)*P*. Substituting PtdIns(4)*P* with the more negatively charged PtdIns(4,5)*P*₂ (10%) also drove exclusive binding through the positively charged surface (**Figure 3C**).

Other negatively charged lipids, phosphatidylserine and cardiolipin (20%), did not replicate this effect (**Figure S3D**), indicating that total negative charge alone is insufficient to promote this alternative membrane association mode. Instead, the specific charge distribution of the inositol headgroup appears critical.

In summary, CG MD simulations suggest that while PtdIns(4)*P* is not strictly required for GOLPH3’s initial membrane association, it controls the binding orientation in a concentration dependent manner.

This novel orientation of GOLPH3 observed in our simulations led us to investigate whether other mechanisms could lead to localization to the *trans*-Golgi. A particularly suited modification is S-acylation, the only reversible modification, which would be compatible with reversible, alternating membrane binding. The SwissPalm webserver (*30*) predicts S-acylation at C122, a cysteine conserved in mouse GOLPH3 but absent in its paralog GOLPH3L and the yeast homolog Vps74p (**Figure S4A**).

To test whether GOLPH3 can be S-acylated, we employed the Acyl-Resin-Assisted Capture (Acyl-RAC) method (*31*), which selectively captures S-acylated proteins by hydrolyzing thioester bonds and binding the resulting free thiols to thiopropyl beads for isolation followed by western blot detection. This approach in HEK293 and HeLa cells indicated that a significant population of endogenous GOLPH3 is S-acylated at steady state (**Figure 3D**). Since Acyl-RAC does not resolve the number of acylated sites, we used Acyl-PEG, a related method where free thiols resulting from hydroxylamine treatment are labelled by PEG-maleimide, which causes gel-shifts proportional to the number of modified cysteines. PEGylation revealed at least three S-acylated cysteines, with little unshifted band, suggesting that the vast majority of GOLPH3 molecules contain at least one acylated site under our experimental conditions (**Figure 3D**).

To confirm S-acylation and identify the sites, we generated five GOLPH3 single-point mutants (Cys-to-Ala), along with a quintuple mutant (GOLPH3-5CA) in which all cysteines were mutated. Using [³H]-palmitate metabolic labeling (*32*) (see Methods), we found that wild-type GOLPH3 is robustly palmitoylated. Mutation of C84 markedly reduced labeling (∼80% loss), while C108 and C122 mutations had a modest effect (**Figure 3E**). GOLPH3-5CA showed no detectable palmitoylation, indicating that C84, C108, and C122 are *bona fide* palmitoylation sites.

Structural analysis revealed that concurrent acylation at C84, C108, and C122 is incompatible with a single membrane orientation (**Figure S4B**). C84 aligns with the membrane-binding surface that accommodates β-hairpin insertion and PtdIns(4)*P* binding (**Figure 3G** and **S4B**). In contrast, C108 and C122 are positioned on the opposite side of the protein, supporting an alternative orientation involving GOLPH3’s positively charged surface (**Figure 3A)**.

To identify the responsible S-acyltransferases, we used six siRNA pools covering the 23 human ZDHHC enzymes in HeLa cells and assessed GOLPH3 acylation by Acyl-RAC (**Figure S4C-E**). Mixes 1 and 3, which respectively contained ZDHHCs 1, 3, 7, 13, and 17 and ZDHHCs 5, 8, 9, and 20, led to a major decrease in recovery by Acyl-Rac (**Figure S4E)**. Silencing of individual enzymes present in these pools showed that knockdown of ZDHHC5, ZDHHC13, and to a lesser extent ZDHHC3, reduced GOLPH3 acylation. Further dissection using the S-acylation defective mutants revealed that ZDHHC13 targets C84, while ZDHHC5 modifies C108 and C122 (**Figure S4F, G**). Notably, ZDHHC13 localizes to the Golgi, while ZDHHC5 is distributed throughout the secretory pathway and the plasma membrane (*33*, *34*).

We next asked whether acylation affects GOLPH3 localization. Quantitative immunofluorescence showed that mutation of C84 reduced Golgi association, and resulted in GOLPH3 relocation at the cell surface (**Figure 3F**). Conversely, mutation of C108 and/or C122 increased Golgi localization, suggesting that acylation at these sites may antagonize Golgi targeting (**Figure 3F**).

Altogether, these findings support a model in which GOLPH3 binds distinct intracellular membranes via two opposing surfaces. At the Golgi, where ZDHHC13 is localized, GOLPH3 inserts its β-hairpin, interacts with PtdIns(4)*P*, and undergoes acylation at Cys-84 (**Figure 3G, H**). In contrast, at the plasma membrane, it may interact with PtdIns(4,5)*P*₂ through its positively charged surface, encounter ZDHHC5, resulting in modification at Cys-108 and Cys-122, which favors this alternative membrane-binding mode. This model suggests that S-acylation dynamically regulates both the subcellular localization and membrane orientation of GOLPH3.

### Identification of the GOLPH3-client enzyme interacting surface

Given the aforementioned effects of S-acylation on GOLPH3 orientation, and presumably its function, we next investigated which region of the GOLPH3 protein interacts with the cytosolic tails of its client enzymes (*17*). To this end, we employed Nuclear Magnetic Resonance (NMR) spectroscopy, selecting the cytosolic tail of LCS, which exhibits micromolar-range binding affinity for GOLPH3 (*17*) (**Figure 4**).

**Figure 4.**
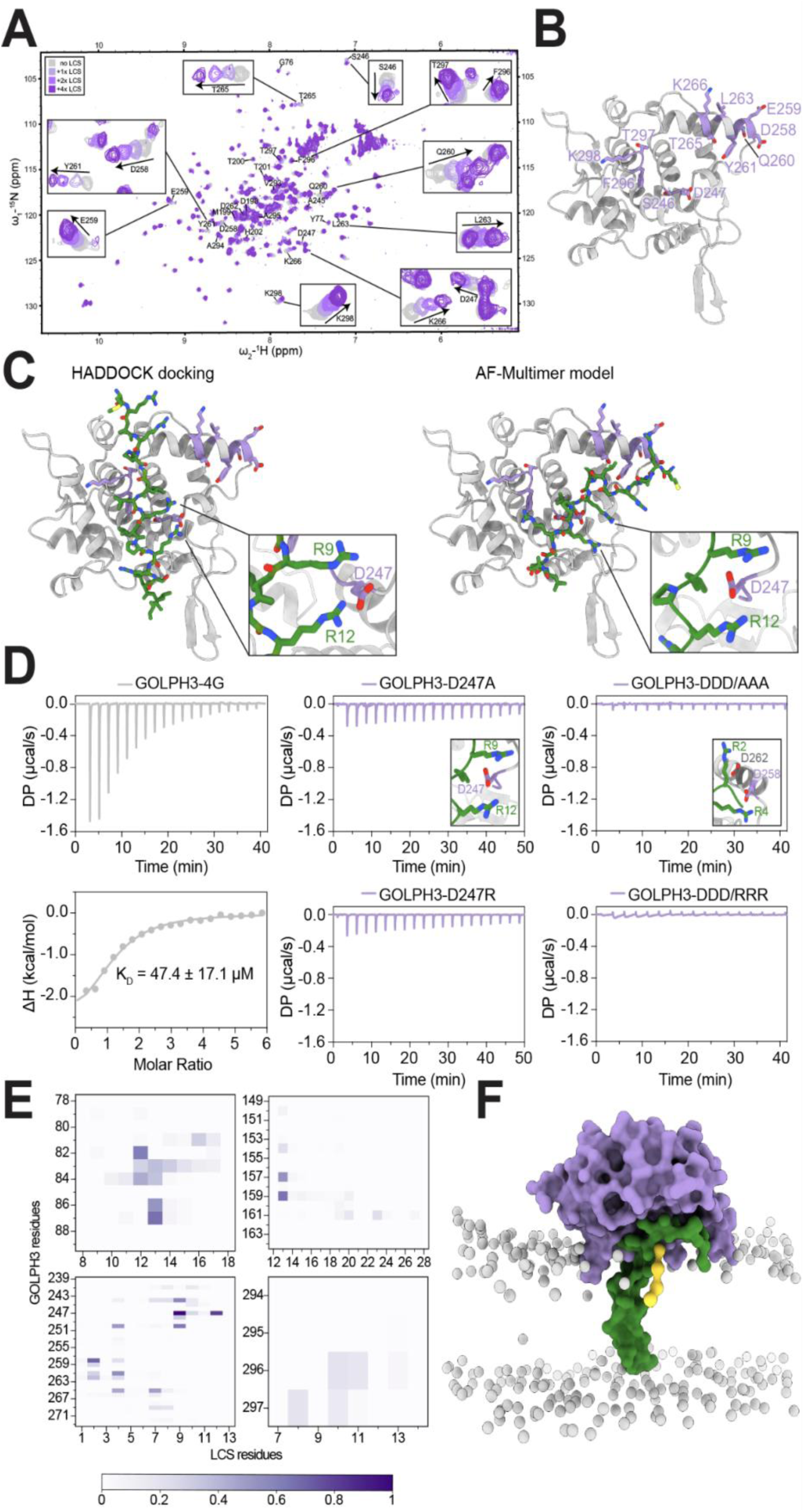
LCS binding site on GOLPH3 as mapped by NMR. A. Overlay of 2D ^1^H–^15^N HSQC spectra of ^15^N-labeled GOLPH3 in its apo form (grey) and with increasing molar ratios of LCS peptide (1:1 to 4:1, gradient grey to violet). Assigned residues are labeled; boxes highlight regions with notable chemical shift changes. **B.** NMR-mapped LCS binding site on the negatively charged surface of GOLPH3, involving residues S246, D247, D258, E259, Q260, Y261, L263, T265, K266, F296, T297, and K298. **C.** Structural models of the GOLPH3–LCS complex from HADDOCK (guided by NMR data) and AlphaFold-Multimer (unguided). Insets show electrostatic interactions. LCS is in green; GOLPH3 residues with NMR shifts are in purple, others in grey. **D.** ITC binding curves for LCS WT with GOLPH3 D247R (left) and D247R/D258R/D262R (middle), and LCS R9A/R12A mutant with GOLPH3 WT (right). Representative of 2–3 replicates. **E.** Normalized contact heatmaps showing residue-level interactions between GOLPH3 and LCS during CG MD simulations. **F.** CG MD snapshot of S-acylated GOLPH3 (at C84) bound to the cytosolic tail and transmembrane domain of LCS (green) at the membrane.

Due to the low solubility of recombinant wild-type (WT) GOLPH3, we engineered a mutant (F194G/L195G/L196G/F197G; hereafter GOLPH3-4G) that reduces hydrophobicity at the β-hairpin tip, improving solubility. GOLPH3-4G exhibited slightly higher thermal stability (**Figure S5A**), remained monomeric by SEC-MALS (**Figure S5B**), retained comparable LCS-binding affinity as WT by isothermal titration calorimetry (ITC) (K_D_ = 65.3 ± 8.9 μM for WT and K_D_ = 47.4 ± 17.1 μM, for GOLPH3-4G; **Figure S5C**), and maintained native folding as shown by overlapping ^1^H, ^15^N HSQC spectra (**Figure S6**).

Titrating the LCS tail peptide into ^15^N-labeled GOLPH3-4G revealed widespread chemical shift perturbations (**Figure 4A**), indicating extensive interaction. The observed fast exchange regime and absence of saturation are consistent with a high micromolar dissociation constant, in agreement with ITC measurements (**Figure S5C**).

To map the binding interface, we assigned perturbed residues using triple-resonance spectra of ^15^N,^13^C-labeled GOLPH3-4G under titration conditions. Despite limited spectral quality, we confidently assigned seven residues that shifted upon peptide binding (S246, D247, T265, K266, F296, T297, K298), and 19 that remained unchanged (**Table S1**, see Methods). Additional assignments via single-residue mutants identified six residues in the 258–263 stretch, five of which (D258, E259, Q260, Y261, L263) shifted upon titration (**Figure S7**).

These data identify a broad, negatively charged interaction surface on GOLPH3, comprising at least S246, D247, D258, E259, Q260, Y261, L263, T265, K266, F296, T297, and K298 (**Figure 4B**). Given the highly basic sequence of the LCS peptide (MRARRGLLRLPRSLLA), this supports a weak, low-specificity electrostatic interaction in solution. However, the effective affinity of the GOLPH3–LCS complex may be higher under physiological conditions, due to the two-dimensional confinement imposed by the membrane surface.

To further explore the interface, we performed computational docking of the LCS peptide to GOLPH3 using HADDOCK (*35*), guided by NMR-identified “active residues”. Docking revealed dominant electrostatic interactions: LCS R2 and R4 with GOLPH3 D273, R9 and R12 with D247, and R13 with E159 (**Figure 4C**). Hydrophobic contacts also contributed, with LCS leucines engaging pockets on GOLPH3: Y241, H244, L272, and V293 with L7; F296 with L8; and W81, S86, and L16 with LCS’s C-terminal leucines. Notably, W81 is a PtdIns(4)*P*-binding residue and may be unavailable in a physiological context.

Because the chemical shift perturbations span a larger area than can be explained by a single docking pose, we used AlphaFold-Multimer (*36*) for blind prediction of the LCS-GOLPH3 complex. The resulting model recapitulated key electrostatic interactions observed in HADDOCK: R2 and R4 with D258 and D262; R9 and R12 with D247; R13 with E159. Hydrophobic interactions were also consistent, involving H244, L249, F253, V268, L272, and Y261 engaging L7–L8 of LCS (**Figure 4C**). Despite not being trained on experimental data, the AlphaFold-Multimer model closely aligned with NMR-mapped residues, reinforcing its validity.

Among the predicted contacts, the interaction between GOLPH3 D247 and LCS R9/R12 emerged as a central binding hub. To validate this, we tested WT GOLPH3 with LCS mutants (R9A, R12A, R9A/R12A) and GOLPH3 mutants (D247A, D247R) using ITC. All mutations reduced binding, with combined mutations of D247, D258, and D262 abolishing interaction (**Figures 4D, S5E**). Importantly, all mutants retained proper folding and thermal stability (**Figure S5F**), confirming that binding loss was not due to protein misfolding.

We next modeled the GOLPH3–client complex in the membrane context using CG MD simulations. The system included the cytosolic tail and transmembrane domain of LCS bound to GOLPH3 on a Golgi-like lipid bilayer, using the AlphaFold-Multimer pose as the starting configuration (**Figure 5C**). Simulations showed stable GOLPH3–LCS association, with the LCS tail exploring the GOLPH3 surface while its transmembrane helix remained embedded (**Figure 5E**).

**Figure 5.**
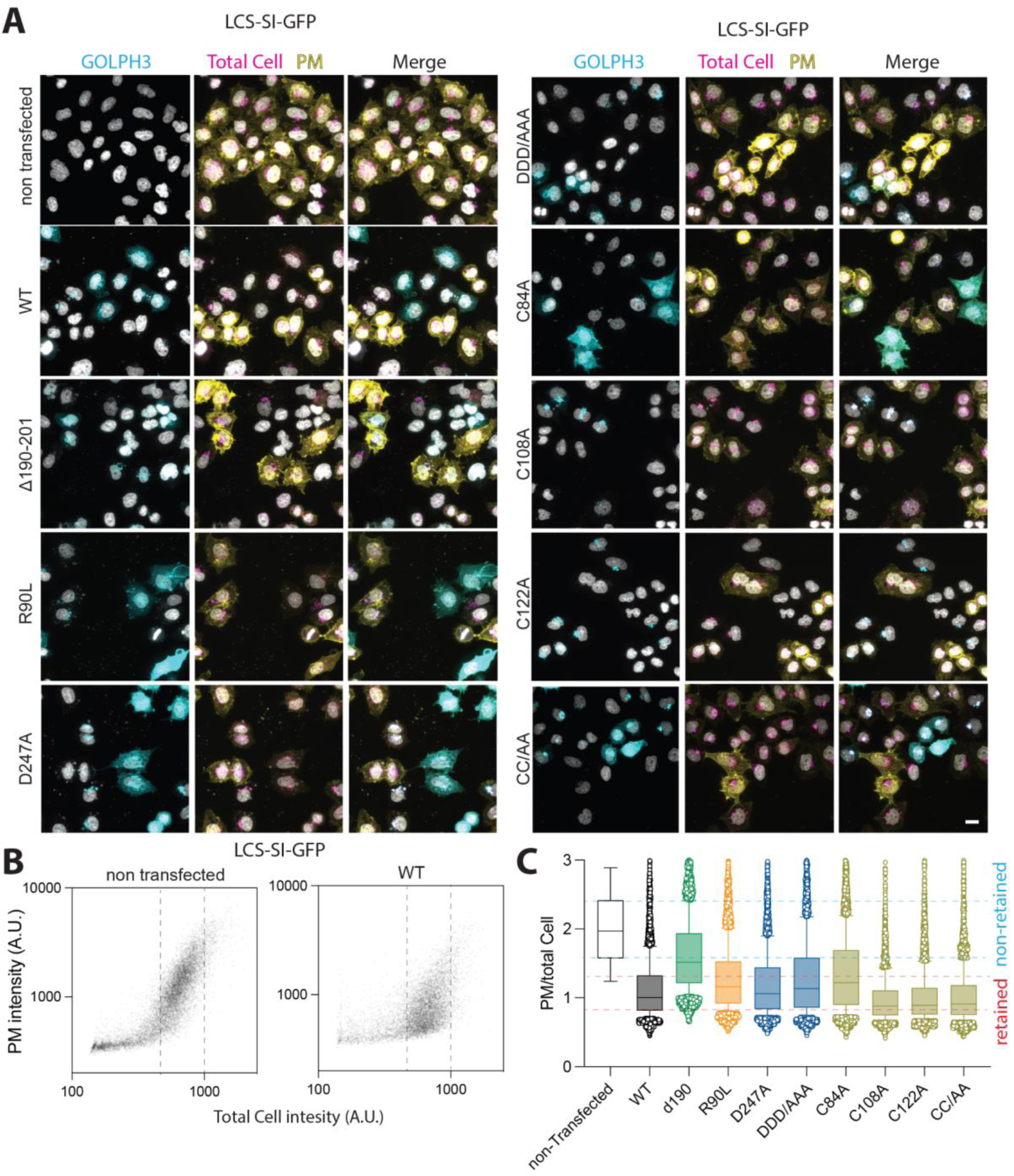
Regulated GOLPH3 interaction with Golgi membranes and clients drives GOLPH3-clients retention at the Golgi. A. Quantitative immunofluorescence of LCS-SI-GFP Golgi retention in GOLPH3 KO HeLa cells transfected with various GOLPH3 variants. Scale bar: 10 µm. Staining: anti-GOLPH3 (cyan), anti-GFP before permeabilization (yellow), LCS-SI-GFP (magenta). **B.** Scatterplot showing total LCS-SI-GFP fluorescence (Total Cell intensity) and surface-exposed signal (PM intensity). Cells were classified as transfected or non-transfected based on GOLPH3 signal. Dashed lines indicate the intensity range used for downstream analysis. **C.** Quantification of LCS-SI-GFP Golgi retention in cells expressing different GOLPH3 mutants, as in (B).

Critically, GOLPH3 maintained a membrane orientation consistent with prior membrane-association modeling and stabilized by C84 S-acylation (**Figure 3G**). When C84 S-acylation was modeled explicitly, GOLPH3 embedded ∼5 Å deeper into the membrane and made ∼20 % more contacts with lipid headgroups, without altering its binding to LCS (**Figures 5F, S5G**).

Altogether, the combination of NMR, MD simulations, AlphaFold predictions, and HADDOCK modeling indicates that GOLPH3 binds its target substrate, LCS, through a well-defined groove lined with negatively charged residues, fully consistent with the membrane-associated conformation in which C84 is S-acylated.

### GOLPH3 regulation drives GOLPH3-clients retention at the Golgi

Once bound to GOLPH3, Golgi-resident proteins are packaged into COPI-coated vesicles, enabling their retrograde transport through maturing Golgi *cisternae* (*17*). We thus investigated how perturbing the various GOLPH3 membrane binding modes described in this study would affect the trafficking of GOLPH3 clients. We made use of a previously established reporter assay based on a chimeric construct (*17*), which consists of the transmembrane domain of the type II plasma membrane enzyme sucrose isomaltase (SI), harbouring GFP at its C-terminus and the cytosolic tail of LCS at the N-terminus . Under normal conditions, the chimera (LCS–SI–GFP) localizes to the Golgi in a GOLPH3-dependent manner (*17*). To assess the extent of reporter mislocalization to the plasma membrane, we used an anti-GFP antibody on non-permeabilized cells, which selectively detects GFP exposed at the cell surface. This allowed us to quantify the fraction of the reporter that escapes Golgi retention, based on the ratio of surface-exposed to total cellular GFP.

Mutants defective in either PtdIns(4)*P* binding (GOLPH3-R90L), membrane association (GOLPH3-Δ190–201), LCS tail-binding (GOLPH3-D247A and the triple mutant GOLPH3-D247A/D258A/D262A, hereafter referred to as GOLPH3-DDD/AAA), or S-acylation (GOLPH3-C84A, GOLPH3-C108A, GOLPH3-C122A, and the double mutant GOLPH3-C108A/C122A) were transfected into GOLPH3-knockout cells. Mutants that impaired GOLPH3 recruitment to the Golgi, particularly the β-hairpin deletion mutant GOLPH3-Δ190– 201, showed markedly reduced retention of the LCS–SI–GFP chimera compared to GOLPH3-WT. Mutations that disrupted LCS binding also diminished retention, although incompletely. The residual activity suggests that, within the two-dimensional context of the membrane, the broad interaction surface between GOLPH3 and its clients is sufficient to maintain partial incorporation into COPI vesicles even when binding is weakened (**Figure 5A, B**).

Interestingly, the effects of S-acylation mutants on LCS retention were divergent. GOLPH3-C84A, which exhibits defective Golgi localization, led to a reduction in retention activity. In contrast, GOLPH3-C108A, GOLPH3-C122A, and the double mutant GOLPH3-C108A/C122A fully retained the reporter at the Golgi, indicating that these modifications do not impair, and may even promote client retention (**Figure 5A, B**).

Together, these findings show that effective binding of GOLPH3 to its client proteins requires membrane association and specific orientation. In this membrane-bound state, the interaction with LCS spans an extended surface of GOLPH3, such that mutations disrupting binding, while sufficient to abolish interaction in biophysical assays, still allow partial functionality in cells. Moreover, S-acylation regulates two functionally distinct states of GOLPH3: acylation at C84 promotes stable Golgi membrane binding and client retention, whereas acylation at C108 and C122 counteracts Golgi localization and opposes LCS retention.

## Discussion

Glycosylation is a non-templated polymerization process (*3*) in which the final glycan structures are largely determined by the sequence of glycosyltransferase encounters with cargoes. Given that the cellular glycome varies across cell states, types, and tissues, a central question in the field is how glycosylation programs are coordinated. Specifically, to what extent is transcriptional control of glycogenes sufficient, and do post-transcriptional mechanisms play an equally critical role? This is particularly relevant in light of our recent findings showing that glycolipids actively shape cell transcriptomes (*37*, *38*).

The Golgi apparatus plays a central role in glycosylation, organizing the spatial and temporal sequence of enzymatic modifications (*1*). This organization depends on the active retention of glycosyltransferases via retrograde COPI-coated vesicles. GOLPH3 is a key COPI adaptor that anchors a specific subset of Golgi enzymes by binding their cytosolic tails (*39*). Modulating GOLPH3 activity, therefore, presents a potential mechanism to selectively reprogram the glycome. This raises critical questions: Are COPI adaptors like GOLPH3 themselves subject to regulation, and what are the consequences of their dysfunction?

In this study, we elucidate the molecular logic underlying GOLPH3 regulation, demonstrating that its membrane association is controlled by specific phosphoinositide interactions and alternative S-acylation events. These modifications toggle GOLPH3 between an active, Golgi-localized, client-binding–competent state and an inactive, plasma membrane–bound state with inverted topology. Notably, we show that GOLPH3 is essential for normal development in mice: its loss disrupts glycosylation and leads to either embryonic lethality or impaired bone mineralization. Interestingly, the Human Genetic Evidence Calculator (HuGE) identifies a very strong association between GOLPH3 and height variation in the human population through rare variant associations (*40*), suggesting potential human correlates of the phenotypes observed in our study.

Our findings reveal a new layer of control in glycosylation: regulated intra-Golgi trafficking that tunes the retention or release of specific glycosyltransferases possibly to reshape the glycome during differentiation. Disruption of this mechanism, as seen in GOLPH3 overexpression in cancer (*23*), highlights its potential role in disease and opens new possibilities for targeted glycome engineering.

## Authors Contribution

A.T. performed the biochemistry and structural biology experiments and contributed to the writing of the manuscript. A.N. conducted the mouse and cell biology experiments and contributed to manuscript writing. L.A.A. performed the NMR and simulation experiments and contributed to the writing of the manuscript. L.A. carried out the S-acylation experiments. F.T. supported the cell biology experiments and generated stable knockout and recombinant cell lines. M.J.M. supervised the biochemistry and structural biology experiments. M.U.A. performed intracellular localization experiments. O.K. conducted the glycosphingolipidomics experiments. S.Y.V. performed the glycoproteomics and proteomics experiments. A.C. assisted with the µCT experiments. S.H. prepared constructs and provided technical support throughout the project. A.A. performed the untargeted lipidomics. L.A. and C.G. assisted with the mouse experiments and data analysis. F.S.M. participated in cell imaging experiments. L.B. provided the mouse tissues. I.K. and N.G. assisted with the mouse experiments. S.V. contributed to the structural biology experiments. A.S. contributed to the cell biology experiments. A.P. contributed simulations. M.M. supervised the glycosphingolipidomics experiments. O.N. supervised the µCT experiments. K.T.S. supervised and analyzed the glycoproteomics and proteomics experiments. F.G.V.D.G. supervised the S-acylation experiments and contributed to manuscript writing. M.D.P. supervised the project and contributed to manuscript writing. G.D.A. conceived and supervised the project and contributed to manuscript writing.

## Acknowledgements

We thank Dimitri Moreau and Stefania Vossio (ACCESS Geneva) for quantitative microscopy and data analysis. Histology data were generated at, or with the assistance of, the EPFL Histology Core Facility. Animal experiments were conducted at, or with the support of, the Center of PhenoGenomics at EPFL. We thank Anne-Laure Mahul Mellier for her support with the animal studies, and Christine Göpfert for the histopathological analysis. G.D.A. is supported by the Swiss Cancer League (KFS-4999-02-2020), the EPFL institutional fund, the Kristian Gerhard Jebsen Foundation, and the Swiss National Science Foundation (SNSF) (310030_184926). M.D.P. is supported by the SNSF (205321_192371) and the EPFL institutional fund. K.T.S. is supported by a Novo Nordisk Foundation Hallas-Møller Ascending Investigator Grant (NNF0073793), the Carlsberg Foundation (CF21-0453), and a Sapere Aude Research Leader Grant from the Independent Research Fund Denmark (2066-00043B). O.K. and M.M. are supported by the Ministry of Agriculture of the Czech Republic (RO0525). We also acknowledge the support provided by the SNSF Gender Equality Grant (310030_184926/2).

## Materials and Methods

### Mice

The generation of GOLPH3^-/-^ mice was done by CRISPR/Cas-mediated genome engineering, generated in collaboration with Cyagen Biosciences, Santa Clara, CA, USA. Exons 2 and 3 were selected as the target site, and Cas9 and gRNA were co-injected into fertilized eggs of C57BL6/J mice. The pups were genotyped by PCR followed by sequencing analysis. The heterozygous progeny (GOLPH3^+/-^) were crossed and used to generate the experimental model used in this study, which are the full systemic GOLPH3 knockout mice (GOLPH3^-/-^).

Notably, exons 2 and 3 encompass residues 74–158. While the vast majority of mouse GOLPH3 transcripts include these exons, our excision strategy does not prevent the expression of isoforms that skip them. Based on our structural and cellular studies, deletion of residues 74–158 is expected to produce a largely non-functional protein. Furthermore, our proteomics and Western blot analyses indicate the absence of full-length GOLPH3 in liver tissue, along with a >75% reduction in the abundance of GOLPH3 tryptic peptides in GOLPH3^-/-^ mice.

All cohorts were inbred at the EPFL breeding facility, and were maintained in a temperature-controlled environment with a 12-h light/12-h dark cycle and free access to standard chow diet and water according to the Swiss Animal Protection Ordinance (OPAn). Male and female animals were used in this study. Body composition (fat and lean mass) was measured using EchoMRI technology with the EchoMRI™ qNMR system (Echo Medical Systems, Houston, TX, USA), as well as whole skeleton morphology and bone mineral density (BMD) by micro - CT using Quantum Fx CT at the Center of PhenoGenomics (CPG) of the EPFL. Unless stated otherwise, animals were sacrificed at room temperature, by cervical dislocation, and all tissues were collected, either snap frozen, fixed for histological analysis. All animal care and treatment procedures were performed in accordance with the Swiss guidelines and were approved by the Canton of Vaud SCAV (authorization VD 3730.c).

### SDS-PAGE and Western Blotting

Proteins from different tissues were extracted using mammalian protein extraction reagent (MPER, Pierce) supplemented with halt phosphatase and halt protease inhibitors (Pierce) as per the manufacturer’s instructions, separated by gel electrophoresis and analyzed using the antibodies anti-GOLPH3/MIDAS (ab98023, Abcam), anti-Calnexin (MAB3126, Millipore) and anti-α-tubulin (ab18027, Abcam).

### RTqPCR

Tissues were powdered manually in liquid nitrogen. RNA was isolated with Tri-Reagent (T9424, Sigma-Aldrich). After centrifugation to remove debris (12,000 g, 10 min, 4°C), chloroform was used for phase separation. The aqueous phase was recovered and precipitated with isopropanol, and centrifuged (12,000 g, 10 min, 4°C), and then, RNA pellets were washed with 75% ethanol. Pellets were resuspended in Milli-Q water. RNA concentrations were determined using NanoDrop and reverse-transcribed using 1,000 ng of RNA and Superscript II enzyme (18064014, Invitrogen) according to the manufacturer’s instructions. qPCR analysis was performed using SYBR Green detection (04913914001, Roche) on a 7900HT Fast Real-Time PCR System (Applied Biosystems) according to the manufacturer’s instructions. Relative mRNA expression was calculated from the comparative threshold cycle (Ct) values of the gene of interest relative to RS9 mRNA.

The specific primer sequences that were used are as follows:

Golph3 (F′-ATAGAATTCATTATGACCTCGCTGACCCAGCGCA, R′-

ATAGGATCCATTTTACTTGGTGAACGCCGCCACCAT) and Rs9 (F′-

CGGCCCGGGAGCTGTTGACG, R′-CTGCTTGCGGACCCTAATGTGACG).

### Liver Tissue Differential O-glycoproteomics

Approximately 30–50 mg of frozen, pulverized liver tissue (Golph3 WT or KO) was homogenized in 500 µL buffer. Samples were heated at 95 °C for 5 min, sonicated (5 × 5 s), and centrifuged at 21,000 × g for 20 min. Proteins were reduced with dithiothreitol and alkylated with iodoacetamide (Sigma), then digested overnight at 37 °C with trypsin (Roche) at a 1:40 ratio. Digests were acidified with trifluoroacetic acid (TFA) for 1 h at 37 °C to inactivate trypsin and quench RapiGest, and desalted using Sep-Pak C18 1cc columns (Waters).

Peptides were quantified using a colorimetric kit (Thermo Fisher), and 200 µg from each sample was labeled with 1.6 mg TMT sixplex (Thermo Fisher). TMT-labelling of peptide samples was prepared as previously described (*41*). Labeled samples were pooled in equal amounts and treated with 0.2 U/mL neuraminidase from Clostridium perfringens (Sigma) to remove sialic acids. Samples were diluted in 2 mL Tris-HCl (175 mM, pH 7.4), and O-glycopeptides (T and Tn antigens) were enriched via lectin weak affinity chromatography (LWAC) using jacalin-agarose beads (Vector Labs), as described in (*42*). Bound glycopeptides were eluted with 3 × 1.0 M D-galactose and desalted using in-house StageTips (C18/C8, Empore 3M). A 50 µg aliquot of the LWAC flow-through was fractionated for proteomic analysis. Both glycoproteome and proteome samples were analyzed by LC-MS/MS.

### Mass Spectrometry-based proteomics

The LC-MS/MS analysis was performed by using an EASY-nLC 1000 UHPLC (Thermo Scientific) interfaced via a PicoView nanoSpray ion source (New Objectives) to an Orbitrap Fusion mass spectrometer (Thermo Scientific). Nano-LC was operated in a single analytical column setup using PicoFrit Emitters (New Objectives, 75-μm inner diameter) packed in-house with Reprosil-Pure-AQ C18 phase (Dr. Maisch, 1.9-μm particle size, ∼19-cm column length), with a flow rate of 200 nl min^−1^. All samples dissolved in 0.1% formic acid were injected onto the column and eluted in a gradient from 2 to 25% acetonitrile in either 95 (for glycoproteomic samples) or 155 min (for proteomic samples), from 25 to 80% acetonitrile in 10 min, followed by isocratic elution at 80% acetonitrile for 15 min (total elution time 120 or 180 min, respectively). The nanoSpray ion source was operated at 2.1-kV spray voltage and 300 °C heated capillary temperature. A precursor MS1 scan (m/z 350–1,700) of intact peptides was acquired in the Orbitrap at a nominal resolution setting of 120,000. For glycoproteomic samples, the five most abundant multiply charged precursor ions in the MS1 spectrum at a minimum MS1 signal threshold of 50,000 were triggered for sequential Orbitrap HCD MS2 and ETD MS2 (m/z of 100–2,000). MS2 spectra were acquired at a resolution of 50,000 for HCD MS2 and 50,000 for ETD MS2. Activation times were 30 and 200 ms for HCD and ETD fragmentation, respectively; isolation width was 4 mass units, and 1 microscan was collected for each spectrum. Automatic gain control targets were 1,000,000 ions for Orbitrap MS1 and 100,000 for MS2 scans, and the automatic gain control for the fluoranthene ion used for ETD was 300,000. Supplemental activation (20%) of the charge-reduced species was used in the ETD analysis to improve fragmentation. Dynamic exclusion for 60 s was used to prevent repeated analysis of the same components.

For proteomic samples, the 10 most abundant multiply charged precursor ions in the MS1 spectrum at a minimum MS1 signal threshold of 100,000 were triggered for sequential Orbitrap HCD MS2 at a resolution of 60,000. In addition, for some proteomic samples, a synchronous-precursor selection MS3 method was used for quantitative analysis (*43*). Polysiloxane ions at m/z 445.12003 were used as a lock mass in all runs. Raw data have been deposited to the ProteomeXchange Consortium (*44*) via the PRIDE partner repository with the data set identifier PXD010155.

### Mass Spectrometry data analysis

Data processing was performed using Proteome Discoverer version 1.4 software (Thermo Scientific) using Sequest HT Node as described previously (*45*) with minor changes. Briefly, all spectra were initially searched with full cleavage specificity, filtered according to the confidence level (medium, low, and unassigned), and further searched with the semi-specific enzymatic cleavage. In all cases, the precursor mass tolerance was set to 6 ppm and fragment ion mass tolerance to 20 milli-mass units. Carbamidomethylation on cysteine residues was used as a fixed modification. Methionine oxidation and HexNAc attachment to serine, threonine, and tyrosine were used as variable modifications for ETD MS2. All HCD MS2 data were preprocessed as described (*46*) and searched under the same conditions mentioned above using only methionine oxidation as a variable modification.

For the quantitative analysis, only HCD MS2 spectra were used. In the case of ETD MS2 spectra, the group of TMT reporter fragment ions (m/z range of 126–132) the quantitative data were extracted from the adjacent HCD MS2 paired spectrum (the same precursor ions), and later used for quantification. Processing of the TMT MS3 data was performed using Proteome Discoverer version 2.1 software (Thermo Scientific).

All spectra were searched against a concatenated forward/reverse human-specific database (UniProt, January 2013, containing 20,232 canonical entries and another 251 common contaminants) using a target false discovery rate of 1%. False discovery rate was calculated using the target decoy peptide–spectrum match validator node. The resulting list was filtered to include only peptides with glycosylation as a modification.

### LC-MS based Bulk Lipidomics

For lipid extraction, samples were prepared following the MTBE protocol (*47*). Briefly, mouse tissues were resuspended in 100 μL of MS-grade water and 360 μL of methanol (MeOH), then homogenized using a handheld homogenizer. A mixture of internal standards (SPLASH® II LIPIDOMIX®; Avanti Polar Lipids, Inc., Alabama, USA) was added. Subsequently, 1.2 mL of MTBE was added, and the samples were vortexed vigorously at maximum speed for 1 hour at 4 °C.

Phase separation was induced by adding 200 μL of MS-grade water, followed by centrifugation at 1,000 × g for 10 minutes. The upper (organic) phase was collected, and the lower (aqueous) phase was re-extracted with 400 μL of a MTBE/MeOH/H₂O mixture (10:3:1.5, v/v/v). This extraction was repeated once more. The combined upper phases were dried under a nitrogen stream and stored at −80 °C.

Eluents consisted of 5 mM ammonium acetate and 0.1% formic acid in water (solvent A), and in isopropanol/acetonitrile (2:1, v/v) (solvent B). Dried lipid extracts were resuspended in a mixture of 20% LC-MS-grade chloroform/methanol (1:1, v/v) and 80% solvent B. Lipid analyses were performed using a Shimadzu Prominence UFPLC XR system (Tokyo, Japan) equipped with a reversed-phase Accucore C30 column (150 × 2.1 mm, 2.6 μm) and a 20 mm guard column (Thermo Fisher Scientific), coupled to a hybrid Orbitrap Elite mass spectrometer (Thermo Fisher Scientific, Bremen, Germany). The instrument was operated in positive ion mode using a heated electrospray ionization (HESI) source.

Lipid quantification was conducted using Skyline (v. 21.1.0.146; MacCoss Lab, University of Washington, Seattle, USA). A transition list was initially generated using LipidCreator, a free and open-source tool integrated with Skyline, and subsequently modified based on experimental data. The transition list included the lipid molecule name, precursor name, precursor m/z, chemical formula, and adduct. Peak areas were normalized to the corresponding internal standards and expressed as a percentage of the total lipid content (mol%).

### GSL Extraction and Analysis

For lipid extraction, cells were harvested into glass tubes (13 × 100 mm) containing 1.5 mL of methanol (Honeywell, Germany). The solution was homogenized using a sonic probe (approximately 20 strong pulses). Mass spectrometry (MS) standards (Avanti Polar Lipids, Alabaster, AL, USA; and Cayman Chemical Company, Ann Arbor, MI, USA) were then added to the samples prior to total lipid extraction.

Following sonication, 0.75 mL of chloroform (Honeywell, Germany) was added. The solution was briefly homogenized again with the sonic probe and left to extract overnight at room temperature. After extraction, the samples were briefly sonicated once more, placed in a heating box, evaporated under a gentle stream of nitrogen (N₂), and then reconstituted in 400 µL of a methanol/chloroform (1:1) mixture. The samples were subsequently transferred to vials, centrifuged, and analyzed via HPLC-MS/MS.

### Liquid Chromatography Separation and Tandem Mass Spectrometry

Sphingolipid species (excluding gangliosides) were separated by reversed-phase HPLC, as previously described (*48*), using a Dionex Ultimate 3000 system (Thermo Scientific, USA) with a Gemini C18 column, 5 µm, 250 × 4.6 mm (Phenomenex, USA), at a flow rate of 0.7 mL/min. A gradient of mobile phases RA (methanol/water, 60:40) and RB (methanol) was used, starting at a 60:40 ratio, with a total run time of 69 minutes. Formic acid and ammonium formate were used as eluent additives.

Glucosylceramide and galactosylceramide were separated by normal-phase HPLC on a Spherisorb 5 µm Silica column, 2.1 × 250 mm (Waters Corporation, Ireland), using a Dionex Ultimate 3000 system (Thermo Scientific, USA). The flow rate was 0.3 mL/min, with a gradient of mobile phases NA (acetonitrile/methanol, 99:1) and NB (acetonitrile/methanol/water, 40:47:13), and a total run time of 48 minutes. Formic acid and ammonium formate were used as eluent additives.

Gangliosides were separated by reversed-phase HPLC on an ARION® C8 column, 4.6 × 300 mm (Chromservis s.r.o., Czech Republic), at a flow rate of 0.8 mL/min. A gradient of mobile phases RA (water/methanol, 90:10) and RB (methanol/isopropanol, 1:1) was applied, starting at a 61:39 ratio, with a total analysis time of 60 minutes. Ammonium formate was used as the eluent additive.

Tandem mass spectrometry was performed using a hybrid triple quadrupole QTRAP 4500 system (AB Sciex, Canada), operating in positive electrospray ionization (ESI) mode. Drying gas, collision energy, and fragmentor voltage were optimized for each species. The instrument was run in multiple reaction monitoring (MRM) scan mode.

### Bone Microarchitecture

To evaluate bone microarchitecture, a SkyScanner 1276 (Bruker, Belgium) was used with a 0.25mm filter, voltage of 200 kV and a current of 55mA. Samples were wrapped in PBS-soaked paper towels and scanned inside a drinking straw sealed on both ends to avoid drying. Voxel size was set at 10x10x10 μm^3^. Bone microarchitecture was evaluated according to the ASBMR guidelines (*49*) using a custom CTan (Bruker, Belgium) script for automatic segmentation of trabecular bone in the distal femoral and proximal tibia VOIs. To account for the marked differences in bone length between the groups, we set the number of slices and the offset with respect to the anatomical landmarks relative to bone length instead of using an absolute number.

The trabecular VOI for femora was defined as the region spanning 15% of the bone length at 7.5% of the bone length away from the distal metaphysis, which resulted in a VOI of about 200 slices with a 100 slice distance from the distal metaphysis for wild-type animals, and VOI of 150 slices 75 slices away from the metaphysis for the GOLPH3 KO mice. The cortical VOI for the femora spanned 10% of the femoral length centred around the midshaft slice, resulting in a VOI of about 130-140 slices for wild-type animals and 100 slices for GOLPH3 KO mice.

Similarly, the trabecular VOI for tibias was defined as the region spanning 12.5% of the bone length at 2.5% of the bone length away from the proximal metaphysis, which resulted in a VOI of about 200 slices with a 40 slice distance from the distal metaphysis for wild-type animals, and VOI of 150-160 slices 30 slices away from the metaphysis for the GOLPH3 KO mice. The cortical VOI for the tibia spanned 5% of the femoral length centred around the slice 20% of the bone length away from the tibio-fibular junction. This results in VOI of about 80 slices 320 slices away from the anatomical landmark for wild-type mice and 60 slices 250 slices away for GOLPH3 KO animals.

The threshold used to binarize the calcified tissue was 40 on a 0-255 scale for trabecular bone and 110 for cortical. Reconstruction of the scans was performed using NRecon (Bruker, Belgium) and further analysis were performed using CTan (Bruker, Belgium) with the minimum for CS to image conversion set at 0 and maximum set at 0.14.

### Tissue Staining

Serial 4 μm paraffin sections were prepared using a rotary microtome and stained with either standard Hematoxylin and Eosin (H&E) to assess general tissue morphology or Alcian Blue (pH 2.5) to visualize cartilage. All staining procedures were performed using a Tissue -Tek Prisma® Automated Slide Stainer (Sakura).

For H&E staining, tissue sections were deparaffinized in xylene, rehydrated through graded ethanol to distilled water, and stained with Harris Hematoxylin solution (Gill II, PAP1; Bio-Optica) for 5 minutes. After rinsing in tap water, sections were briefly differentiated in 1% acid-alcohol (700 mL absolute ethanol, 10 mL 37% HCl, 290 mL H₂O), followed by a 10-minute wash in running tap water for nuclear blueing. Cytoplasmic structures were stained with a working solution of Eosin-Phloxine for 1 minute, prepared by combining 50 mL of 1% Eosin Y (Sigma, E4382), 5 mL of 1% Phloxine B (Fluka, 28550), 360 mL of 95% ethanol, and 2 mL of glacial acetic acid. Slides were then washed in distilled water, dehydrated, cleared in xylene, and mounted using Eukitt mounting medium (Sigma-Aldrich).

For Alcian Blue staining, sections were deparaffinized, rehydrated, and incubated for 25 minutes at room temperature in 1% Alcian Blue (Sigma-Aldrich, A5268) dissolved in 3% acetic acid (pH 2.5). The solution was filtered prior to use and prepared by dissolving 5 g of Alcian Blue in 500 mL of 3% glacial acetic acid. After staining, slides were washed in running tap water for 5 minutes and briefly rinsed in distilled water. Nuclei were counterstained with 0.1% nuclear fast red (Vector Laboratories, H-3403) for 5 minutes, followed by rinsing, dehydration, clearing, and mounting with Eukitt. Acidic mucopolysaccharides and mucins stained blue, while nuclei appeared red.

### Molecular Dynamics simulations

All MD simulations were set up with the CHARMM-GUI server (*50*, *51*) produced with Gromacs 2022 or 2024, and analyzed in VMD assisted with custom scripts. Within CHARMM-GUI, coarse-grained systems were parametrized with the MARTINI 2.2p (*52*) forcefield starting from the GOLPH3 crystal structure PDB 3KN1 (*12*).

For CG simulations aimed at probing spontaneous binding of GOLPH3 to various membranes, the protein was initially positioned such that its closest bead was 28-30 Å from the nearest membrane bead. The membrane was composed of a mixture of approximately 30% POPC, 20% DOPC, 12% POPE, 7% DOPE, 5% POPS, and 16% cholesterol and a variable lipid component consisting of either 0%, 5%, 10%, 20%, or 40% PtdIns(4)*P*, or 10% PtdIns(4,5)*P*_2_, CL, or PS, as explained for each simulation and in each case proportionally reducing the fractions of the first set of lipids. The membrane was parameterized with standard MARTINI 2 lipids, and the system was solvated with the polarizable water and 150 mM NaCl. Each system was minimized and equilibrated with the standard CHARMM-GUI protocol, that is by slowly removing restraints in the NVT ensemble using Particle Mesh Ewald for the electrostatic contributions and velocity rescale for temperature coupling up to 310 K. The production phase was carried out in the NPT ensemble using a Parrinello-Rahman semiisotropic coupling algorithm for maintaining the pressure constant at 1 bar, 310 K as the target temperature, and 20 fs integration time steps. Five independent replicas were run for all simulations.

For CG simulations of GOLPH3 S-acylated at C84 on the membrane, we added the 4 beads that represent the palmitate group in the MARTINI 2.2p framework extending away from the C84’s sidechain bead, and applied the parameters optimized in (*53*).

For CG simulations of the GOLPH3 – LCS complex on the membrane, we took the NMR-consistent model of GOLPH3 and LCS generated with AlphaFold-Multimer and extended the LCS towards its C-terminus with a canonical helix of the corresponding amino acid sequence, projecting into the membrane. The parametrization of this system proceeded as explained above, including the same procedure to model the acylated form of C84; and the system was minimized, equilibrated and produced in MD with the same protocols.

### Generation of stable cell lines expressing LCS-SI-GFP

The LCS-SI-GFP plasmid was previously used and described in (*17*). HeLa cells were transfected with the plasmid using JetPrime transfection reagents (Polyplus) following the manufacturer’s instructions. Stable clones were selected using 800 μg/mL of G418 antibiotic. Expression of the GFP-tagged proteins in the selected clones was confirmed by Western blot analysis using an anti-GFP antibody (Ref 11814460001, Roche).

### Generation of GOLPH3 KO/LCS-SI-GFP clones

GOLPH3 KO cell line was generated from stable LCS-SI-GFP HeLa cells using CRISPR/Cas9 technology. These cells were transfected with GOLPH3 CRISPR plasmids (sc-412973, Santa Cruz Biotechnology) by using JetPrime transfection reagents (Polyplus) following the manufacturer’s instructions. Single-cell clones were isolated by limiting dilution in 96-well plates. The resulting clones were screened for GOLPH3 KO by Western blot analysis using an anti-GOLPH3 antibody (ab98023, Abcam).

### Acyl-Resin Assisted Capture (Acyl-RAC) assay

Protein S-acylation was assessed by the Acyl-RAC assay as described by (*54*), with some modifications. HeLa or HEK-293 cells were lysed with a buffer containing 0.5% Triton-X100, 100 mM Hepes pH 7.4, PBS, 1 mM EDTA, 0.2 mM SDS and protease inhibitor cocktail). In order to block free SH groups with 1 M N-Ethylmaleimide (NEM), 200 ml of blocking buffer (100 mM Hepes, 1 mM EDTA, 87.5 mM SDS and 50 mM NEM) was added to cell lysate and incubated for 2 hours at 40 °C. Subsequently, 3 volumes of ice-cold 100% acetone was added to the blocking protein mixture and incubated for 20 minutes at 20 °C and then centrifuged at 5,000 g for 10 minutes at 4 °C to pellet precipitated proteins. The pellet was washed five times in 1 ml of 70% (v/v) acetone and resuspended in a buffer (100 mM Hepes, 1 mM EDTA, 35 mM SDS). For treatment with hydroxylamine (HA) and capture by Thiopropyl Sepharose beads, 2 M HA was added together with the beads (previously activated for 15 min with water) to a final concentration of 0.5 M HA and 10% (w/v) beads. As a negative control, 2 M Tris was used instead of HA. These samples were then incubated for 3 hours at room temperature on a rotating wheel. The beads were washed, the proteins were eluted from the beads by incubations in 40 μl SDS sample buffer with beta-mercapto-ethanol for 5 minutes at 95 °C. Finally, samples were submitted to SDS-PAGE and analysed by immunoblotting. The same protocol was used to identify the palmitoyltransferases responsible for GOLPH3 S-acylation through an siRNA screening of ZDHHC enzymes. HeLa cells were transfected with siRNAs against six different mixes of ZDHHC enzymes. Mix 1 included ZDHHC 1, 3, 7, 13, and 17, Mix 2: ZDHHC 2, 4, 6, and 16, Mix 3: ZDHHC 5, 8, 9, and 20, Mix 4: ZDHHC 12, 15, and 18, Mix 5: ZDHHC 11, 23, and 24, and Mix 6: ZDHHC 14, 19, 21, and 22.

### PEGylation assay

The level of protein S-palmitoylation was assessed as described in (*55*), with minor modifications. HeLa or HEK-293 cells were lysed with the following buffer 0.5% Triton-X100, 100 mM Hepes pH 7.4, PBS, 1 mM EDTA, 0.2 mM SDS and protease inhibitor cocktail. After centrifugation at 100,000 g for 15 minutes, supernatant proteins were reduced with 25 mM TCEP for 30 minutes at room temperature, and free cysteine residues were alkylated with 1 M NEM for 2 hours at room temperature to be blocked. After chloroform/methanol precipitation, resuspended proteins in PBS with 4% SDS and 5 mM EDTA were incubated in buffer (1% SDS, 5 mM EDTA, 1 M NH2OH, pH 7.0) for 1 hour at 37°C to cleave palmitoylation thioester bonds. As a negative control, 1 M Tris-HCl, pH 7.0, was used. After precipitation, resuspended proteins in PBS with 4% SDS were PEGylated with 20 mM mPEG-5 for 1 hour at 37 °C to label newly exposed cysteinyl thiols. After precipitation, proteins were resuspended with SDS sample buffer and boiled at 95 °C for 5 minutes.

### Radiolabeling experiments

To detect palmitoylation, HeLa WT or GOLPH3 KO cells were transfected or not with different GOLPH3 constructs, incubated for 3 hours in IM (Glasgow minimal essential medium buffered with 10 mM Hepes, pH 7.4) with 200 mCi/ml ^3^H palmitic acid (9,10-^3^H(N)) (American Radiolabeled Chemicals, Inc.). The cells were washed and directly lysed for immunoprecipitation with the anti-GOLPH3 rabbit antibody. For all radiolabeling experiments, after immunoprecipitation, washed beads were incubated for 5 minutes at 90 °C in reducing sample buffer prior to 4–12% gradient SDS-PAGE. After SDS-PAGE, the gel was incubated in a fixative solution (25% isopropanol, 65% H_2_O, 10% acetic acid), followed by a 30-minute incubation with signal enhancer Amplify NAMP100 (GE Healthcare). The radiolabeled products were revealed using Typhoon phospho-imager and quantified using the Typhoon Imager (ImageQuanTool, GE Healthcare). Quantification of radioactive experiments was quantified using specific screens by autoradiography. The images shown for ^3^H-palmitate labeling were however obtained using fluorography (indirect detection of radioactive emission by stimulated light emission from a fluorophore (Amplify) on film).

### Immunofluorescence and confocal microscopy

HeLa GOLPH3 KO cells transfected with the different GOLPH3 constructs were plated at approximately 50% confluency post-transfection. Cells were washed with PBS, fixed in 4% paraformaldehyde (PFA) for 20 minutes, washed with PBS, quenched with 50 mM NH_4_Cl for 10 minutes, and washed again with PBS. Cells were permeabilized with 0.05% saponin in PBS for 5 minutes. Subsequently, cells were washed and blocked with 3% BSA and 0.05% saponin in PBS for 30 minutes. Primary antibodies (anti-GOLPH3 and anti-GM130) and secondary antibodies were diluted in the same buffer. After blocking, coverslips were washed and incubated with primary antibodies overnight at 4 °C, washed in PBS, and incubated for 45 minutes with secondary antibodies. Cells were then washed and stained with Hoechst for nuclear visualization. Finally, coverslips were mounted in Prolong Glass Antifade Mountant. Images were acquired using a Zeiss LSM 980 Inverted microscope with a 63X objective, a pinhole size of 1 AU and at least 2 line-averaging. Images were processed using Fiji (ImageJ).

### LCS-SI-GFP retention assay

HeLa GOLPH3 KO/LCS-SI-GFP cells were transfected with the different GOLPH3 constructs. Forty-eight hours after transfection, cells were harvested by trypsinization, fixed 10 minutes in 4% PFA/PBS, washed, and resuspended in PBS containing 1% BSA.

To assess plasma membrane (PM) localization of luminal GFP chimeras containing the transmembrane domain of sucrase–isomaltase and the corresponding N-terminal LCS cytosolic tails, cells were incubated with 4 μg/ml anti-GFP mouse monoclonal antibody (Ref. 11814460001, Roche) and 4 μg/ml of anti-mouse Alexa Fluor 568 (A-10037, Thermo Fisher Scientific) for 1 hour at RT without permeabilization.

After incubation, cells were extensively washed with PBS/1% BSA, fixed 10 minutes in 4% PFA/PBS again and washed. For selective detection of cells expressing GOLPH3 constructs, a rabbit polyclonal anti-GOLPH3 antibody (2 μg/ml; Abcam, ab98023) and an anti-rabbit Alexa Fluor 647 (2 μg/ml; A-31573 Thermo Fischer Scientific) were applied in permeabilization conditions (PBS/1% BSA and 0,05% Saponin) for 1 hour at RT. Cells were then washed and stained with Hoechst for nuclear staining.

Negative controls included cells stained with secondary antibodies only and unstained cells. Data acquisition was performed using an LSRII SORP flow cytometer (BD Biosciences).

### Bacterial protein expression and purification

The human GOLPH3 (Uniprot sequence Q9H4A6, residues 52-298) expression plasmid carries the gene for the GOLPH3 protein lacking the first 51 amino acids, with an N-terminal 10xHistidine tag and a Twin-Strep-tag, followed by a TEV (Tobacco Etch Virus) cleavage site. GOLPH3 was cloned into the bacterial expression vector pET_HST and mutations were introduced using the GenScript’s Site-Directed Mutagenesis Services (GenScript). The coding sequences for all GOLPH3 variants, except for the GOLPH3 WT and GOLPH3 D247A mutants, were optimized for Escherichia coli (*E. coli*) expression (Genscript). The plasmids were transformed into BL21(DE3) competent cells. Protein expression was induced with 0.5 mM isopropyl b-D-1-thiogalactopyranoside (IPTG) when cells reached an optical density (OD at 600 nm) of 0.6, with a subsequent growth overnight at 20 °C. Cell pellets were resuspended in lysis buffer (500 mM NaCl, 50 mM Tris pH 8.0, 1 mM TCEP, 10% glycerol) and were lysed by sonication. After centrifugation at 20,000 xg for 60 minutes at 4 °C, the supernatant was filtered and applied to a HisTrap excel column (Cytiva) at room temperature. After elution of recombinant GOLPH3 with a continuous gradient over 30 column volumes of elution buffer (500 mM NaCl, 50 mM Tris pH 8.0, 1 mM TCEP, 10% glycerol, 500 mM imidazole), pure fractions were buffer exchanged into final buffer (250 mM NaCl, 50 mM Tris pH 8.0, 1 mM TCEP) to remove imidazole, using a HiPrep Desalting column (GE Healthcare). The N-terminal purification tags were cleaved by overnight incubation with TEV protease at room temperature. Subsequently, the cleaved tags and TEV protease were removed with another His affinity chromatography and the flowthrough was collected, concentrated, and additionally purified by Size Exclusion Chromatography (HiLoad 16/600 Superdex 200 pg, Cytiva) in final buffer. Pooled fractions were concentrated to the required concentration using a MWCO concentrator (Millipore) with a cutoff of 10 kDa.

### Size exclusion chromatography coupled to multi-angle light scattering

The molecular weights of the tested constructs were determined by size exclusion chromatography coupled to multi-angle light scattering (SEC-MALS). The mass measurements were performed on a UltiMate3000 HPLC system equipped with a 3 angles miniDAWN TREOS static light scattering detector (Wyatt Technology). The sample volumes of 5−10 μl at a concentration of 12 mg/mL, were applied to a Superdex 200 5/150 GL column (Cytiva) previously equilibrated with 250 mM NaCl, 50 mM Tris pH 8.0, 1 mM TCEP at a flow rate of 0.08 mL/min. The data were analyzed using the ASTRA 6.1 software package (Wyatt technology), using the absorbance at 280 nm and the theoretical extinction coefficient for concentration measurements.

### Nuclear Magnetic Resonance (NMR) Spectroscopy

All proteins used for NMR experiments were designed on the GOLPH3-4G background due to its higher stability and solubility. Recombinant proteins were prepared with the isotope enrichment scheme appropriate for each experiment (i.e. ^15^N-only or ^15^N, ^13^C) in 20 mM MES buffer pH 6.5 with 250 mM NaCl and 10% ^2^H_2_O. Lower salt, even by 10%, yielded much degraded spectra. Protein concentrations were around 300 µM for all NMR experiments, limited by protein stability and solubility.

All NMR experiments were carried out in a 18.8 T (800 MHz ^1^H Larmor frequency) Bruker spectrometer equipped with a CPTC ^1^H, ^13^C, ^15^N 5 mm cryoprobe and an Avance Neo console. The temperature was set to 308 K in all experiments, as spectra were substantially better than at the more standard 298 K yet without compromising the limited stability. Backbone (H, N, CA, and C) and CB resonances were assigned using a standard procedure based on conventional 3D HNCA, HN(CO)CA, HNCO, HN(CA)CO, CBCA(CO)NH and HNCACB spectra. All spectra were acquired and processed using Bruker’s Topspin 4.0 software. Backbone assignments were obtained through manual spectral analysis assisted by the program CARA. As explained, the protein assignment is only partial, and assisted by HSQC spectra acquired on single-point mutants. The NMR chemical shifts for all confidently assigned residues are provided in **Table S1**.

### Isothermal titration calorimetry

Isothermal titration calorimetry (ITC) experiments were performed on a MicroCal PEAQ-ITC (Malvern Panalytical), with GOLPH3 variants being in a buffer containing 250 mM NaCl, 50 mM Tris pH 8.0 and 1 mM TCEP. LCS peptides were synthesised and delivered as lyophilised powder (Genscript). The peptides were dissolved in the protein buffer and their concentrations were calculated based upon their absorbance at 280 nm and their corresponding molar extinction coefficient. Experiments consisted of titrations of a single 0.4 µL injection followed by 19 injections of 2 μl of LCS peptides, at approximately 2.5 mM, into the cell containing GOLPH3 proteins at a 100 μM concentration. Experiments were performed at 25 °C in triplicate (n = 3), with injection duration of 4 s, injection spacing of 150 s, stir speed of 750 rpm, and reference power of 7 μcal/s. Data were processed and plotted using MicroCal PEAQ-ITC Analysis Software and fitted to a one-site binding model.

### Circular Dichroism Spectroscopy

Circular Dichroism Spectra of GOLPH3 proteins (at 10 μM in 100 mM NaCl, 50 mM Tris pH 7.4 and 1 mM TCEP) were collected with a Chirascan V100 (AppliedPhotophysics), using a 1 mm path length cuvette, at 20 °C, in the 200-260 nm range. Thermal denaturation experiments were performed by heating the sample from 20 to 95 °C and collecting CD data at 207 nm every 1 °C, with a temperature ramp speed of 1 °C/min and an equilibration time of 15 seconds.

---

**Figure S1:**
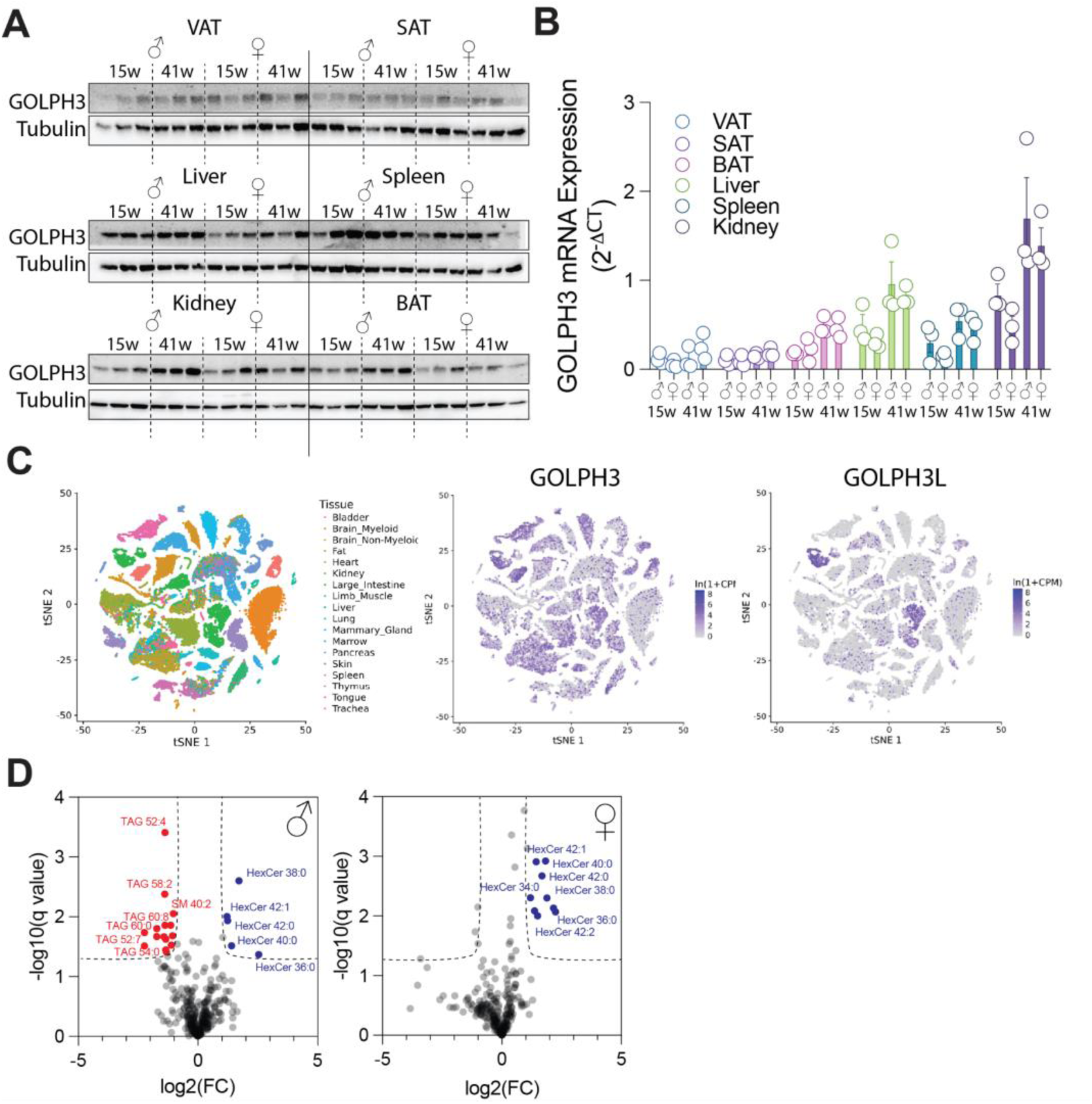
GOLPH3 KO causes glycosylation defects *in vivo*. A. Western blot analysis of lysates from visceral adipose tissue (VAT), subcutaneous adipose tissue (SAT), liver, spleen, kidney, and brown adipose tissue (BAT) from young (15 weeks) and old (41 weeks) male and female mice. GOLPH3 and tubulin (loading control) protein levels were detected using specific antibodies. **B.** Quantitative PCR (qPCR) analysis of GOLPH3 mRNA levels in VAT, SAT, liver, spleen, kidney, and BAT from the same mouse cohorts. **C.** t-SNE plot derived from the Tabula Muris single-cell RNA sequencing dataset (*24*), showing the expression patterns of GOLPH3 and GOLPH3L across multiple organs and tissues. **D.** Volcano plot illustrating lipid species that differ significantly in abundance between livers of GOLPH3^+/+^ and GOLPH3^-/-^ mice (n = 5 per group), based on liquid chromatography–mass spectrometry (LC-MS)–based lipidomics. Lipid species reduced in GOLPH3^-/-^ livers are shown in red; those increased are shown in blue.

**Figure S2:**
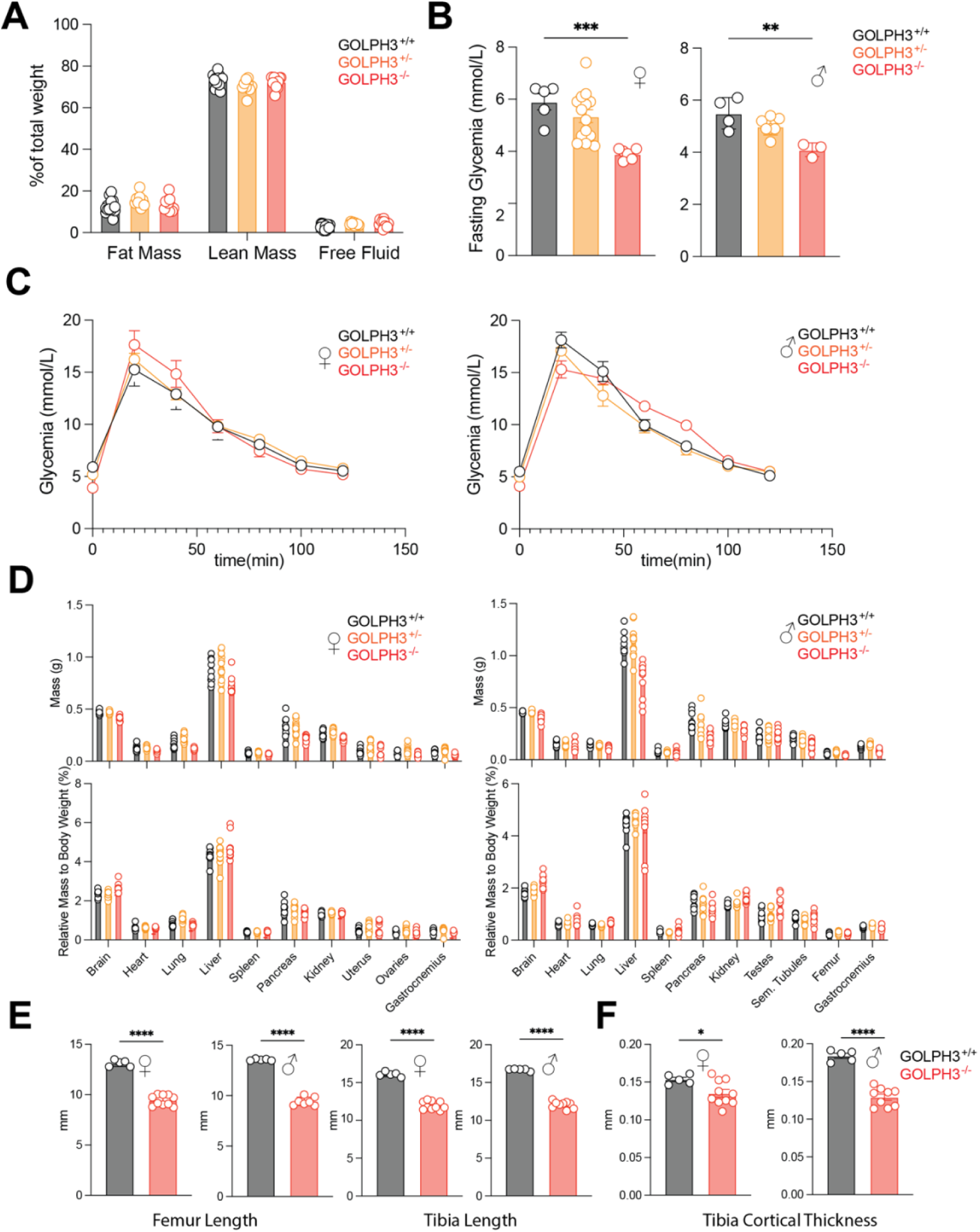
Metabolic and skeletal phenotyping of GOLPH3-deficient mice. A. Histogram showing body composition in 9-week-old GOLPH3^+/+^, GOLPH3^+/-^, and GOLPH3^-/-^ mice, as determined by Echo-MRI. Data reflect the percentage of total body weight attributable to fat and lean mass. Values represent mean ± SEM. **B.** Fasting blood glucose levels in 9-month-old GOLPH3^+/+^, GOLPH3^+/-^, and GOLPH3^-/-^ mice. Data are presented as mean ± SEM (two-way ANOVA, **p < 0.001; ***p < 0.001). **C.** Blood glucose concentrations measured over a 2-hour period following glucose administration in 9-month-old GOLPH3^+/+^, GOLPH3^+/-^, and GOLPH3^-/-^ mice. Scatter plot shows mean ± SEM. **D.** Histograms displaying the absolute (top) and relative (bottom) weights of internal organs in female (left) and male (right) GOLPH3^+/+^, GOLPH3^+/-^, and GOLPH3^-/-^ mice. Data represent mean ± SEM. **E.** Bone length measurements of the femur and tibia in 9-week-old male and female GOLPH3^+/+^ and GOLPH3^-/-^ mice. Data are shown as mean ± SEM (unpaired t-test, ****p<0.0001). **F.** Tibial cortical thickness in 9-week-old male and female GOLPH3^+/+^ and GOLPH3^-/-^ mice, quantified by µCT analysis. Values represent mean ± SEM (unpaired t-test, *p < 0.05; ****p<0.0001).

**Figure S3.**
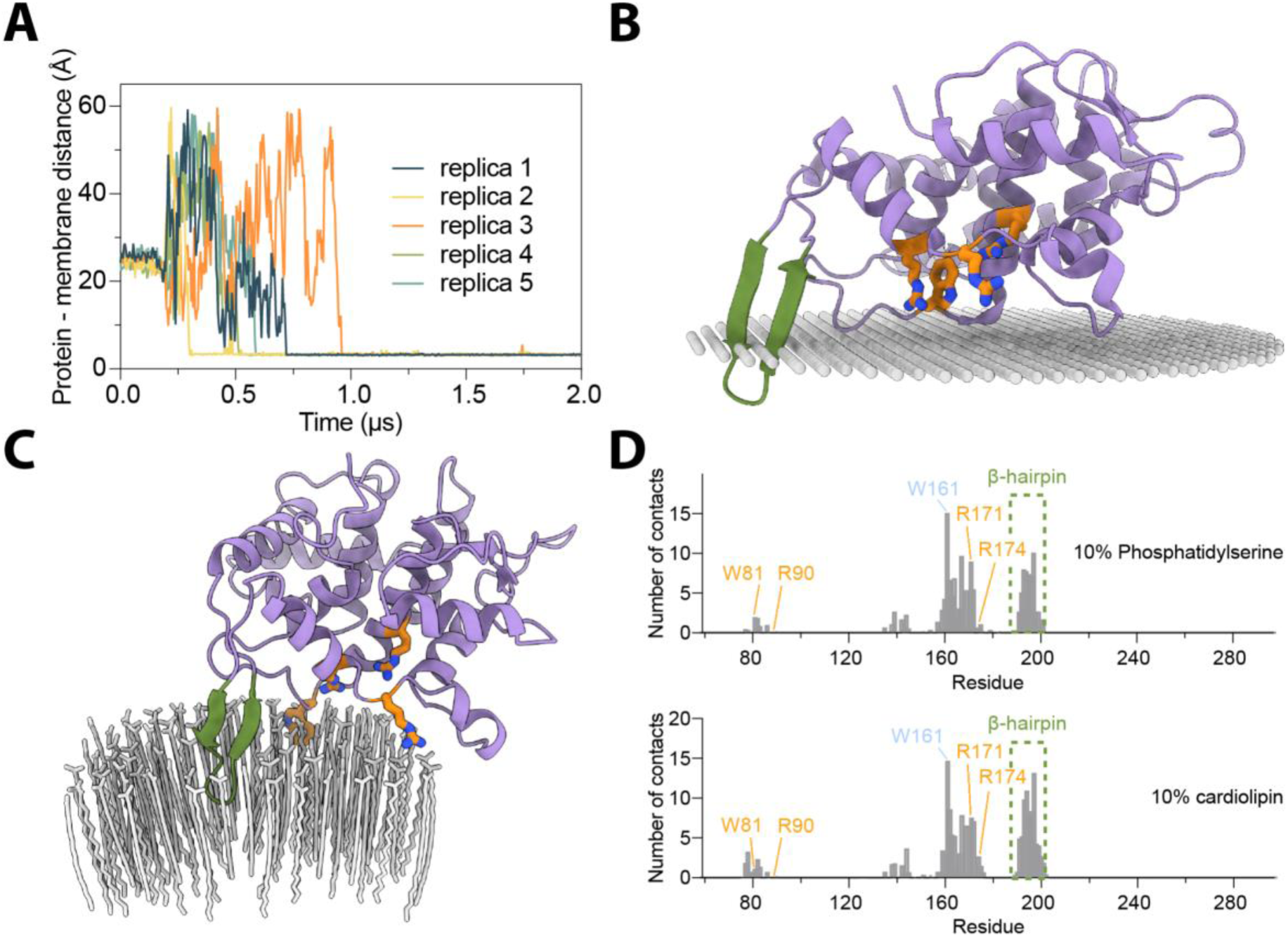
Predicted membrane orientation of GOLPH3. A. Minimum distance over time between any coarse-grained (CG) bead of GOLPH3 and the closest membrane bead across five independent CG-MD simulations with membranes containing 5% PtdIns(4)*P*. **B.** Predicted orientation of GOLPH3 on the membrane from the OPM server. Hydrophobic boundaries are shown as grey spheres; the β-hairpin is in green, and PtdIns(4)*P*-binding residues are shown as orange sticks. **C.** GOLPH3 orientation relative to membrane mimics (oleic, palmitic, and myristic acids, grey sticks) predicted by AlphaFold3, using the maximum number of lipid molecules allowed. **D.** Average number of contacts per residue between GOLPH3 and membranes containing 10% phosphatidylserine (top) or 10% cardiolipin (bottom), based on CG MD trajectories.

**Figure S4.**
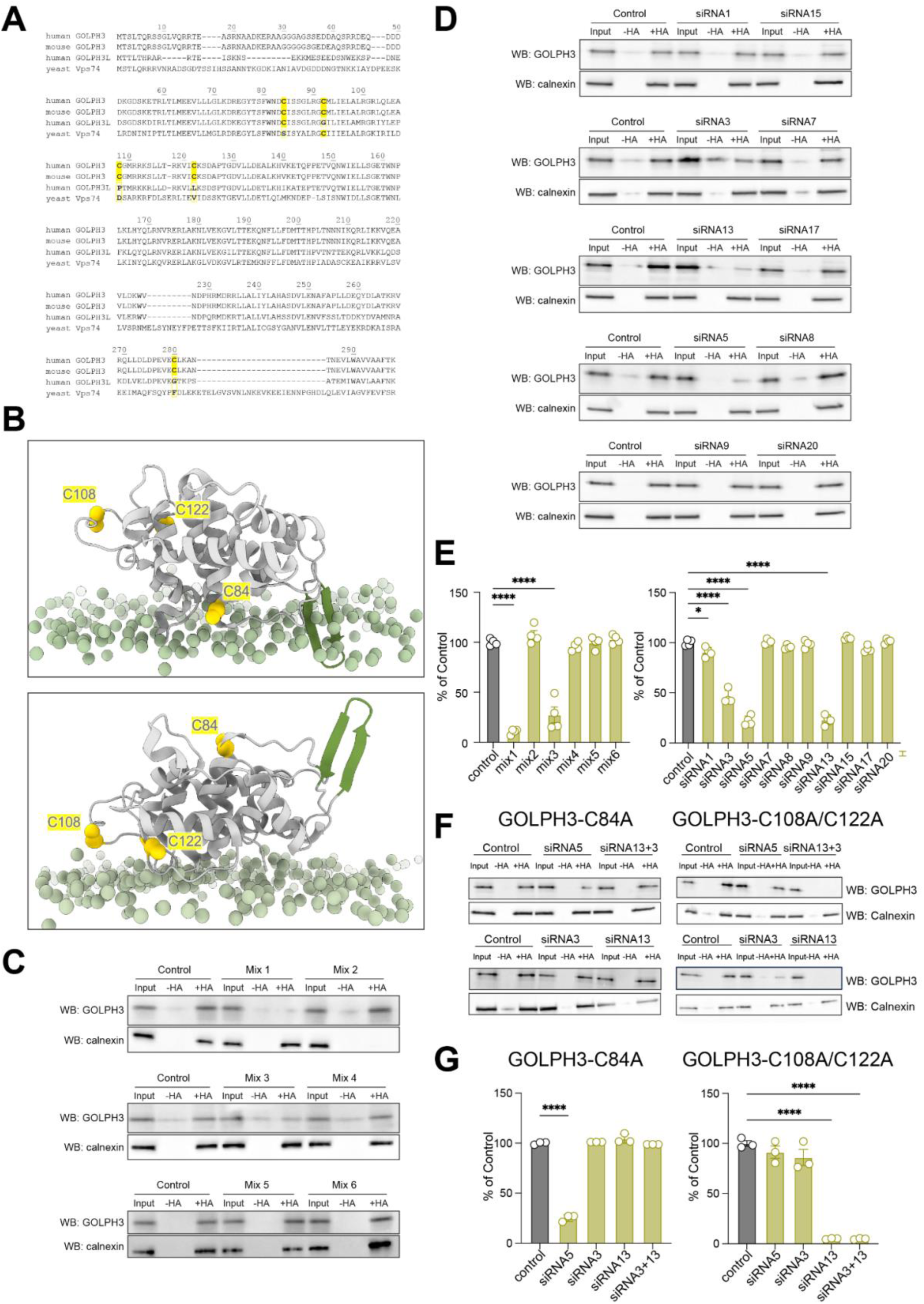
GOLPH3 undergoes palmitoylation on Cys84, Cys108 and Cys122. A. Sequence alignment of human GOLPH3, mouse GOLPH3, human GOLPH3L, and yeast Vps74, highlighting the five cysteine residues in human GOLPH3 (yellow). Alignment was performed using Clustal Omega. **B.** Positions of the three palmitoylated cysteines on GOLPH3 relative to the membrane, shown in two MD-derived orientations: via the β-hairpin (top) and via the positively charged surface (bottom). **C–D.** S-acylation of endogenous GOLPH3 in HeLa cells following siRNA knockdown of ZDHHC enzymes (C: siRNA mixes; D: individual siRNAs). Palmitoylation detected after hydroxylamine treatment (+HA); immunoblotting with anti-GOLPH3. Calnexin served as loading control. **E.** Quantification of endogenous GOLPH3 S-acylation from (C–D). Mean ± SEM, n = 4. One-way ANOVA vs. control: : *p < 0.05; ****p < 0.0001. **F.** S-acylation of GOLPH3-C84A and GOLPH3-C108A/C122A mutants in GOLPH3 KO HeLa cells treated with ZDHHC-targeting siRNAs. **G.** Quantification of mutant GOLPH3 S-acylation from (F). Mean ± SEM, n = 3. One-way ANOVA vs. control: : ****p < 0.0001.

**Figure S5.**
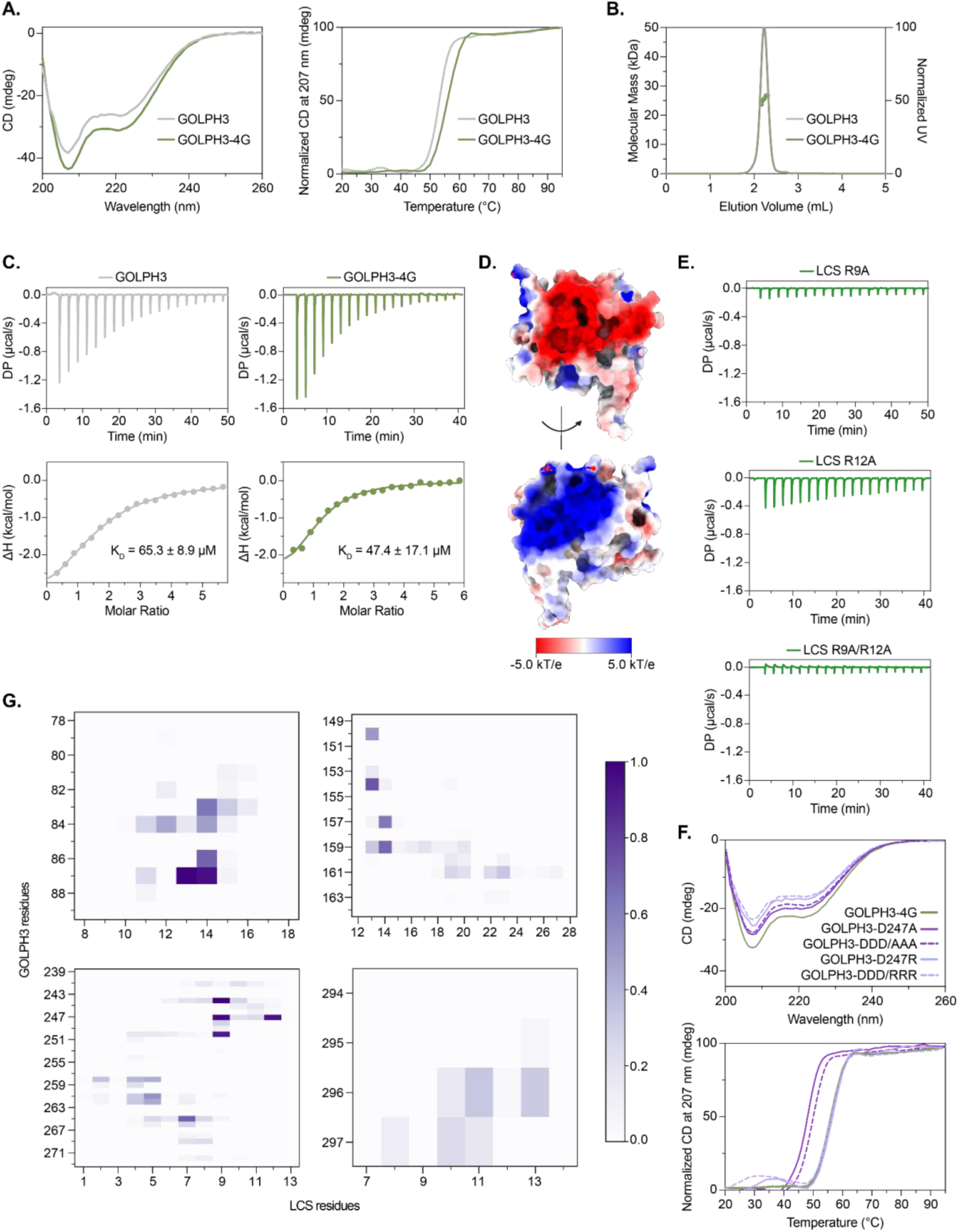
LCS binding site on GOLPH3. A. CD spectra of GOLPH3-4G reveal secondary structure identical to WT. Left: normalized thermal denaturation monitored at 207 nm; Tm = 52.5 °C (WT) and 55 °C (4G). B. SEC-MALS confirms that both GOLPH3 WT and 4G are monomeric. C. ITC measurements of LCS binding to GOLPH3 WT and 4G. Representative traces shown; K_D_ values are means from n = 3 technical replicates. D. Electrostatic surface potential of GOLPH3 WT computed via APBS. E. ITC traces showing binding of LCS mutants R9A (top), R12A (middle), and R9A/R12A (bottom) to GOLPH3 WT (n = 2 replicates). F. CD spectra of GOLPH3 mutants confirm WT-like secondary structure. Right: melting temperatures 4G (55 °C), D247A (48.5 °C), D247R (57 °C), D247A/D258A/D262A (49.5 °C), D247R/D258R/D262R (57 °C). G. Normalized heatmap showing residue-level contacts between S-acylated C84 GOLPH3 and LCS during MD simulations. Scale bar: contact intensity.

**Figure S6.**
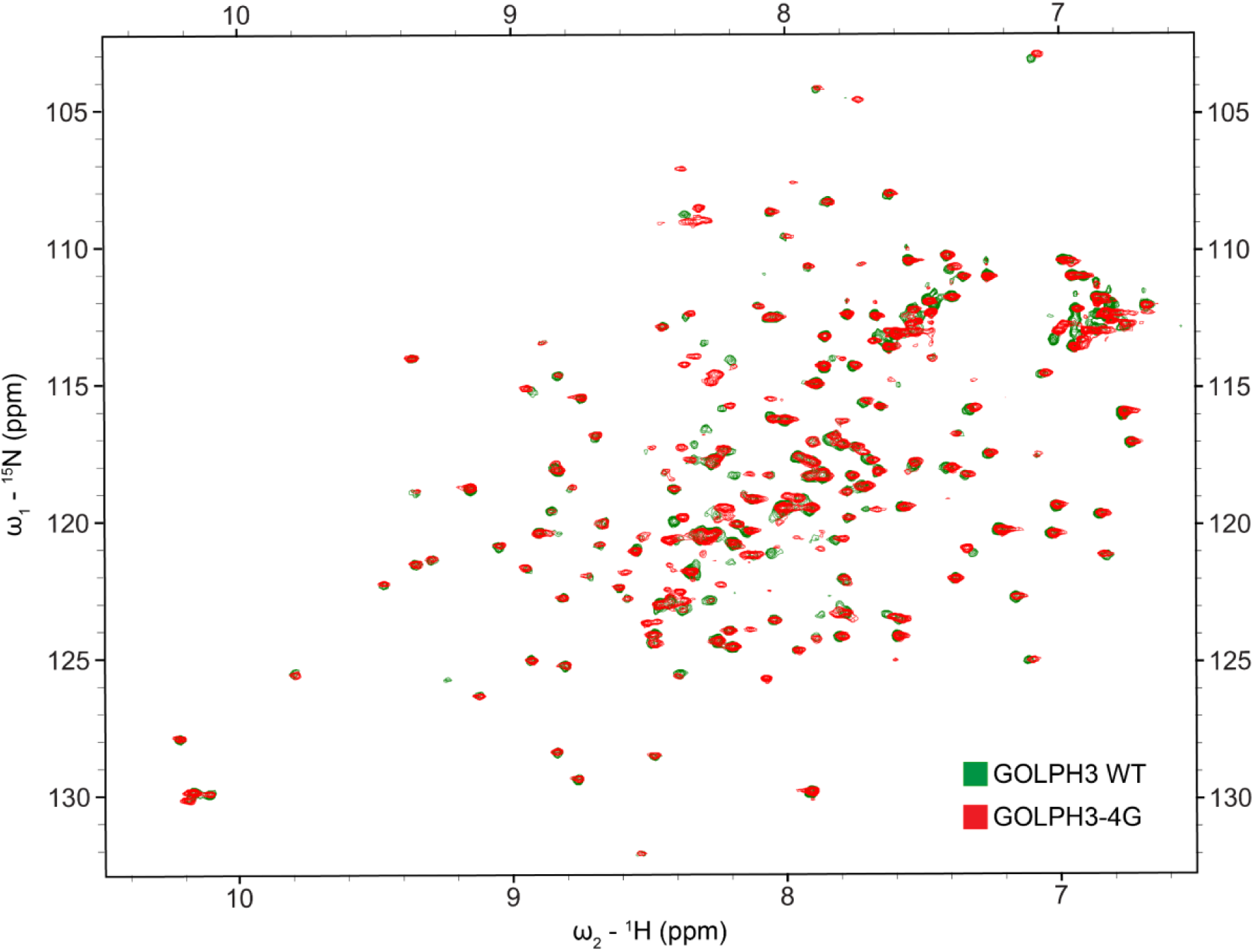
GOLPH3-4G shares the same fold as the WT protein. Overlay of the 2D ^1^H-^15^N HSQC spectra of ^15^N-labeled GOLPH3 WT (green) and ^15^N-labeled GOLPH3-4G mutant (red).

**Figure S7.**
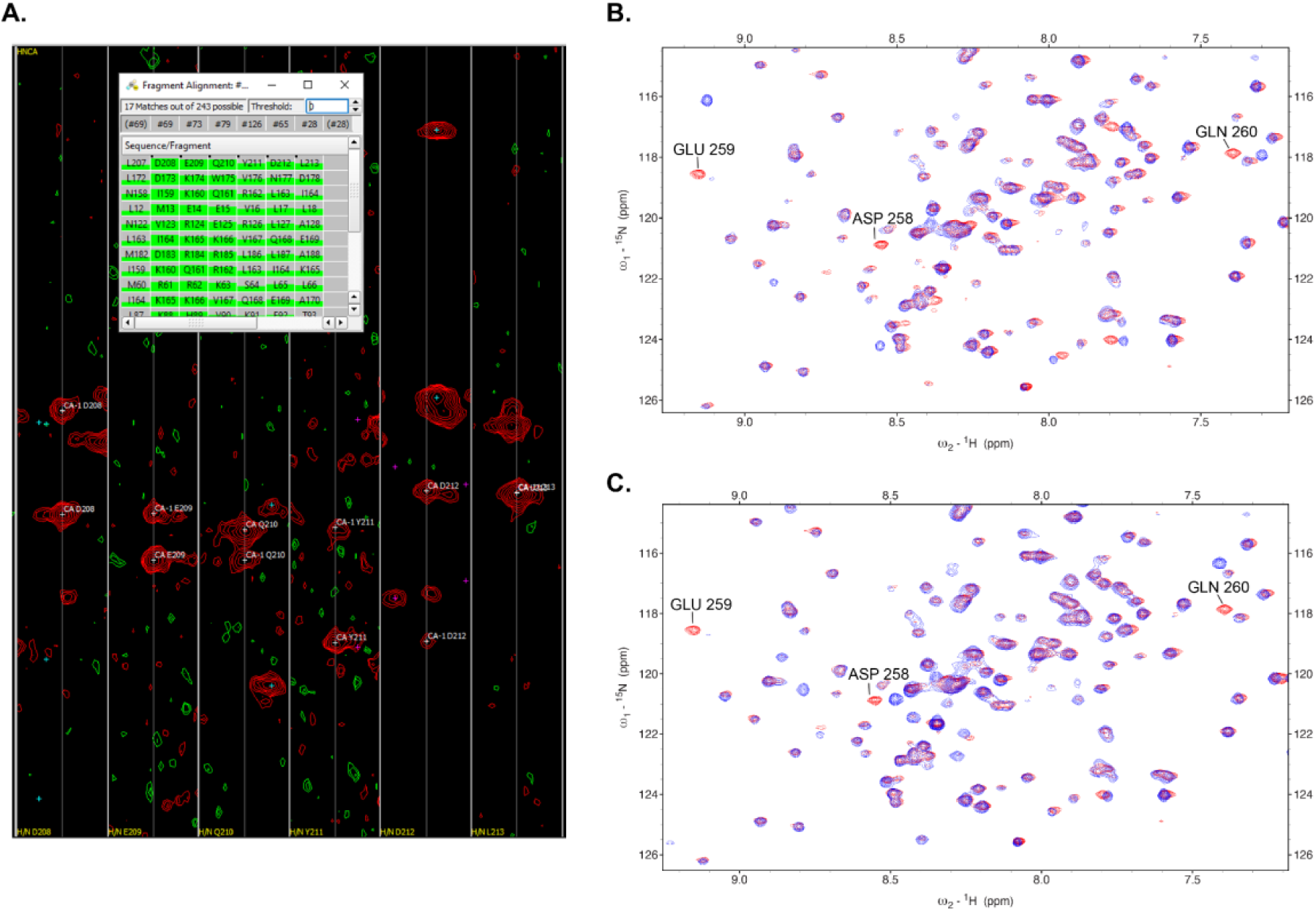
Confirming assignments for GOLPH3-4G’s segment D258 to L263. A. HNCA strips showing a fragment of linked spin systems in CARA, best matching the sequence D_258_EQYDL_263_ (noting a -50 residue offset in the software), though chemical shifts could also fit other segments. **B.** HSQC spectra of GOLPH3-4G (red) and D258A (blue) mutants. Substantial shifts in D258, E259, and Q260 confirm the assignment. **C.** HSQC spectra of GOLPH3-4G (red) and E259A (blue) mutants. Mutations cause clear shifts in E259, D258, and Q260. Together, these confirm the assignment of the D258–L263 segment.

**Table S1.** Backbone chemical shifts for GOLPH3-4G (ppm). As explained in the methods, chemical shifts are in MES pH 6.5 at 308 K.

| Residue | H | N | CA | CB | C |
| --- | --- | --- | --- | --- | --- |
| G76 | 7.7302 | 104.4004 | 43.1271 |  |  |
| Y77 | 7.3368 | 120.7746 | 52.6107 | 39.3925 |  |
| G195 | 8.2864 | 108.7969 | 42.6103 |  |  |
| G196 | 8.3448 | 108.9055 | 42.6175 |  |  |
| G197 | 8.3115 | 108.75 | 42.6223 |  | 171.2667 |
| D198 | 8.2825 | 120.2743 | 51.655 | 38.5638 | 173.8144 |
| M199 | 8.4254 | 120.4332 | 53.0886 | 29.9318 | 173.8309 |
| T200 | 8.2657 | 114.4234 | 59.7786 | 67.1529 | 171.9781 |
| T201 | 8.0514 | 115.3135 | 58.8964 | 67.4606 |  |
| H202 | 8.2315 | 122.0959 | 51.5815 | 27.7911 | 170.2667 |
| A245 | 7.5637 | 119.252 | 51.9491 | 16.8112 | 175.965 |
| S246 | 7.0681 | 102.7262 | 58.198 |  | 171.6638 |
| D247 | 7.6041 | 123.2618 | 51.7653 |  |  |
| D258 | 8.5439 | 120.8278 | 55.6249 | 37.8208 | 174.9889 |
| E259 | 9.1364 | 118.5201 | 57.1687 | 26.2028 | 176.7921 |
| Q260 | 7.3913 | 117.8191 | 56.1395 | 26.6791 | 174.1618 |
| Y261 | 8.7091 | 121.7632 | 59.9624 |  |  |
| D262 | 8.3054 | 120.0985 | 54.8162 |  |  |
| L263 | 7.2088 | 120.1409 | 54.8897 |  |  |
| T265 | 7.6116 | 107.8329 | 63.1971 | 66.048 |  |
| K266 | 7.7896 | 123.9872 | 57.132 | 29.3794 | 176.7756 |
| V293 | 7.949 | 117.3537 | 65.1085 |  |  |
| A294 | 8.5961 | 122.1431 | 52.5004 | 14.8086 | 176.6763 |
| A295 | 8.1231 | 120.1343 | 51.655 | 14.6878 | 176.1139 |
| F296 | 7.7406 | 114.0837 | 56.0292 | 37.0001 | 173.9136 |
| T297 | 7.8556 | 114.0999 | 60.5505 | 67.4982 | 171.1509 |
| K298 | 7.902 | 129.6113 | 54.853 | 30.708 | 178.7442 |

